# Elucidating Partial Folding State of Bovine Pancreatic Trypsin Inhibitor by a Combined Study of Molecular Dynamics Simulations, Information Theory, Molecular Graph Theory, and Machine Learning

**DOI:** 10.1101/2023.11.14.566993

**Authors:** Hiqmet Kamberaj

**Author notes:** Corresponding author *Email address:* (Hiqmet Kamberaj).

## Abstract

Using a notably large amount of data in investigating physical and chemical phenomena demands new statistical and computational approaches; besides, the cross-validations require well-established theoretical frameworks. This study aims to validate the statistical efficiency of alternative definitions for the information-theoretic measures, such as transfer entropy, using the so-called (*α, q*)-framework. The primary goal is to find measurements of high-order correlations that preserve information-theoretic properties of information transfer between the components of a dynamical system (such as a protein) due to local operations. Besides, this study aims to decode the information contained in the amino acid sequence establishing a three-dimensional protein structure by comparing the amino acids physical-chemical properties with their ranked role in the protein interaction network topology using new graph-theoretic measures based on the constructed digraph models of (*α, q*) information transfer within a heat flow kernel embedding framework. Moreover, this study aims to use the Deep Graph Convolution Neural Networks for classifying the role of each amino acid in a protein trained upon short equilibrium structure fluctuations at sub-nanosecond time scales.

In particular, this study examines the influence of disulphide bridges on the three-dimensional structure of the Bovine Pancreatic Trypsin Inhibitor wild type and mutated analogue protein.

## 1. Introduction

Although slow conformational motions of large biological molecules studies are limited in time and size scales, atomistic models (AMs) combined with the molecular dynamics (MD) technique are widely used to understand the internal atomic motion of biomolecular systems [1, 2, 3]. Furthermore, a considerably large amount of the data collected to investigate physical and chemical phenomena will require new statistical and computational approaches to be significantly efficient. Moreover, the cross-validation of these new computational approaches requires vast data collected from the simulations and well-established theoretical frameworks.

The problem of encoding the protein structure into amino acid sequences remains unsolved. Information contained in the amino acid sequences can be classified as overlapping of physical-chemical properties of amino acids. The information-theoretic measures, such as mutual information [4], have already been considered [5] to be sensitive to linear and nonlinear statistical correlations, and an improvement compared to Pearson correlation measures [6], supported by some previous studies [7, 8]. These measures can measure the statistical mutual information content between the amino acids of a protein, considered components of an interaction network system, by focusing on the position fluctuations of a group of atoms. On the other hand, correlation coefficients (such as mutual information or Pearson correlation coefficient) are symmetric measures; that is, from the measured mutual information content, one can not distinguish the *sender* (or *source*) and *receiver* (or *sink*) amino acids of the information flow in the interaction network of amino acid sequences [8]. Therefore, the so-called *transfer entropy* [9, 10, 11, 12, 13] showed as an information-theoretic measure of the information flow. The transfer entropy is also sensitive to nonlinear statistical dependences. It enables, as an asymmetric measure, measuring the directional information flow between two amino acids in a protein interaction network and distinguishing between the source amino acid (that is, the driver amino acid of the interaction network pathway signal) and the sink amino acid (that is, the signal receiver) [8]. The transfer entropy has successfully been used in studying signal pathway sampling in biological system [14], altering the information flow in the amino acid interaction network due to mutation for studying partial protein unfolding [8], changes in the information flow pathways in proteins due to ligand binding [15], studying the hinge influence on the LAC repressor headpiece, information flow in the protein allostery [16, 17], studying allosteric communication landscapes in proteins [18, 19], and studying the heat flow random walks at the protein-RNA interface [20].

The statistical analysis frameworks have been established [20] (and the references therein) to study the amino acids interaction network associated with the topology of the information flow in proteins. The weighted directed graph model was chosen to describe mathematically the topological network of the anisotropic diffusion of the heat flow in proteins. Furthermore, the information-theoretic measures, such as Shannon formalism of transfer entropy [9, 8], are used to describe the geometrical properties of the graphs, defined by the equilibrium structure fluctuations at sub-nanoseconds time scales from molecular dynamics simulations.

In this study, we aim to cross-validate the statistical efficiency of newly developed alternative definitions for the information-theoretic measures of the transfer entropy between time series, based on the so-called (*α, q*)-framework introduced in Ref. [21]. The primary goal is to find measurements of high-order correlations that satisfy information-theoretic properties of information transfer, mainly due to local operations [22]. Besides, they should be easy to compute using the existing algorithms [8, 23]. Furthermore, this study aims to decode the information contained in the amino acid sequence that establishes a protein structure by comparing the physical-chemical properties of amino acids with their role in the protein interaction network topology; for that, new graph-theoretic measures are introduced based on the constructed digraph models of (*α, q*) information transfer between amino acids [20]. The primary aim is to develop a data-driven approach based on causal reconstruction and graph network measures on the causal effects [24] that will identify amino acids acting as major system structure drivers and gateways (or hubs) of equilibrium structure fluctuations at a given time scale from MD simulations.

Moreover, this study aims to use the so-called Deep Graph Convolution Neural Networks (DGCNN) [25], using a new algorithm developed in this study, for classifying the amino acid role in a protein structure trained upon short equilibrium structure fluctuations at sub-nanoseconds time scales from molecular dynamics simulations. In this study, a heat kernel embedding framework [26, 27] is employed to encapsulate information flow in a DiGraph using eigenvalue decomposition of the Laplacian to compute the embedding coordinates matrix of every vertex in the graph.

In particular, this study aims to examine the influence of disulphide bridges on the three-dimensional (3D) structure of the Bovine Pancreatic Trypsin Inhibitor (BPTI) protein (wild type and its mutated analogue) [28, 29].

## 2. The (*α, q*)-Framework of Information Flow

In the Supporting Information materials associated with this study, we introduce the so-called (*α, q*)-framework, according to Ref. [21]. In particular, we describe the (*α, q*)-definitions of the entropy and mutual information for the discrete probability distribution Markovian time processes, aiming at maximising the mutual information and hence the channel capacity. Then, we have extended formalism for the continuous probability distribution Markovian time processes, by defining the Sharma-Mittal [30], Rényi [31], and Tsallis [32] differential entropies and corresponding mutual information. Besides, we cross-validate the statistical efficiency of (*α, q*)-framework by comparing it with Shannon formalism for different values of *α* and *q*; in this study, we consider two different intuitive linearly and non-linearly coupled Markovian processes describing the input and output of a Gaussian-like noisy channel and XOR Bernoulli-like noisy channel, respectively. Moreover, we describe the numerical analysis of two newly developed algorithms for computation of the information-theoretic measures, based on the so-called symbolic encoding and kernel density estimators using artificial intelligence techniques provided in Python programming language paradigm within *sklearn* package. Note that the possibilities for extending the Shannon framework of the information theory have already been attempted [33], relying on parametrisation of the information content by introducing the non-additivity property in uncertainty. The advantage is that these new frameworks allow developing higher-order measures sensitive to the non-linear couplings and noisy signals in dynamical systems.

In the following, we introduce a new theoretical framework for the transfer entropy, the so-called (*α, q*) transfer entropy.

### 2.1. The (α, q) Transfer Entropy

The transfer entropy, *T*_*Y* → *X*_, is defined as [9]:

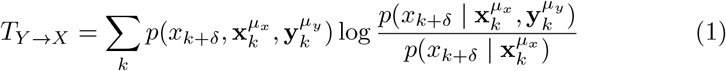

where *k* is a time index and the information is in units of *nats*. The vectors 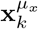 and 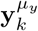 are the state vectors of the Markovian processes and *μ*_*x*/*y*_ ≡ (*m*_*x/y*_, *τ*_*x/y*_) are the embedded parameters, as described in Supporting information. Similarly, the transfer entropy, *T*_*X* → *Y*_, is given as:

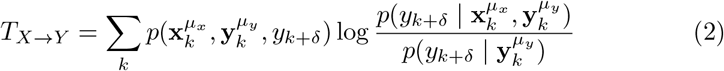

This formulation of the transfer entropy represents a dynamical measure, as a generalisation of the entropy rate to more than one element to form a mutual information rate [9]. In Eq. 1, 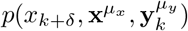 is the joint probability distribution of observing future value *x*_k+ *δ*_ and the pasts of 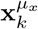 and 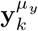 is the conditional probability of observing the future value of *X* given the past values of both *X* and *Y*, and 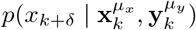 is the conditional probability of observing the future of *X* knowing its past. From Eq. 1, the transfer entropy can also be expressed in terms of Shannon entropies:

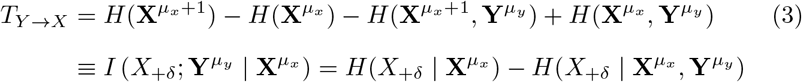

where 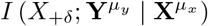 is the conditional mutual information [11, 8, 34]. Equation 3 indicates that the transfer entropy, *T*_*Y* → *X*_, can be interpreted as the average amount of information contained in the source about next state *X*_+*δ*_ of the variable *X* that was not already contained in the past of *X*.

Similarly, *T*_*X* → *Y*_ is given as

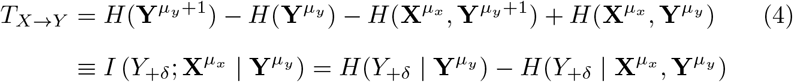

In this study, the statistical efficiency of alternative definitions of the transfer entropy measures between time series is validated, based on (*α, q*)-framework [21], as described in Supporting Information. Note that the volume of an *d*-dimensional space is such that 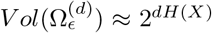, which is the volume of the smallest set 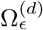 that contains most of probability distribution for a random process *X* [4]. That is, in *d*-dimensional, small Shannon entropy values imply that the random process is confined to a small effective volume while large entropy values imply that the random process is confined to a large effective volume. By definition [4], the set 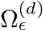 is the smallest volume set with probability 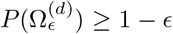 to the first order in the exponent, where ϵ is an infinitesimally small number. That corresponds to Gaussian-like Markovian processes, which are characterised by short tails, and hence these measures do not maximise the mutual information or channel capacity, in particular, for noisy channels. In contrary, (*α, q*)-information measures can be used to count for noisy random processes characterised by longer tails. In particular, the primary goal is to find measurements of high-order correlations that satisfy information-theoretic properties of information transfer, mainly due to local operations [22], and being easy to compute using the existing algorithms [8, 23].

In this study, we define new measures of the transfer entropies, the so-called here (*α, q*) transfer entropies, as:

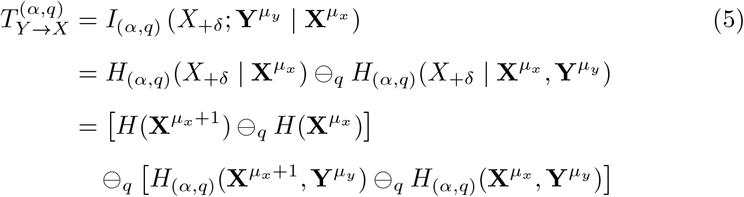

Similarly,

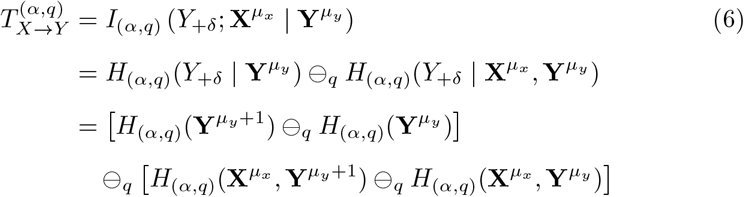

Here, the function *x* ⊖_*q*_ *y* is defined as [21]:

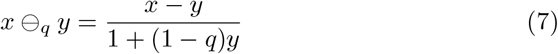

which reduces to standard difference for *q* = 1. In Eq. 5 (or Eq. 6), in general, *H*_(*α*,q)_(*X*) is the Sharma-Mittal entropy [30]:

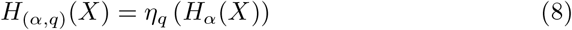

Here, *H*_*α*_(*X*) is Rényi entropy (see Eq. SI2 in Supporting Information) and η_q_ is the function [21]:

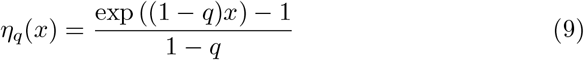

Note that for *q* = 1 and *α* ≠ 1, equations 5 and 6 reduce to Rényi transfer entropy formulations [35], based on *α*-framework of entropy while for *q* ≠ 1 and *α* = 1, they reduce to the so-called here Tsallis transfer entropies, based on the so-called here *q*-framework. When *q* = *α* = 1, equations 5 and 6 give the standard transfer entropies based in the Shannon entropy formulation [9].

### 2.2. Tsallis Local Transfer Entropy

The so-called local Shannon transfer entropy measure is defined as [36]:

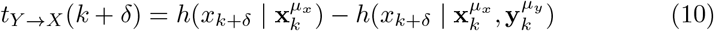

Note that everywhere mentioned in this study, *δ* denotes a time parameter, which indicates an average time-length correlation between *X* and *Y* (typically, *δ* = 1). In the right-hand side of Eq. 10, the local condition entropies are defined as:

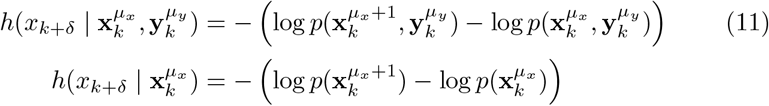

Similarly, the local transfer entropy, *t*_*X* → *Y*_ (*k* + *δ*), is defined as:

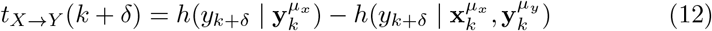

where

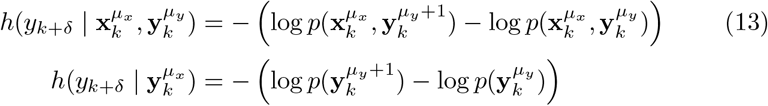

In this study, we introduce a new measure of the local transfer entropy, named as *Tsallis local transfer entropy*, based on the Tsallis *q*-framework:

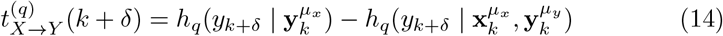

where

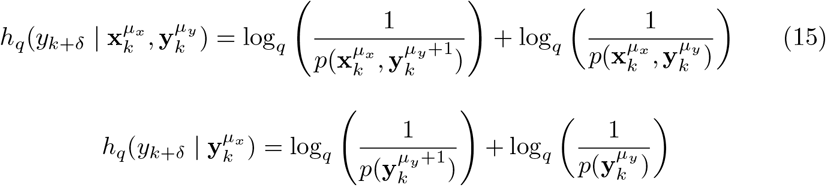

In Eq. 15, the function log_q_(*x*) is defined as [32, 37, 38]:

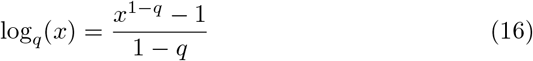

Note that using Eq. 12 and Eq. 14, we can have:

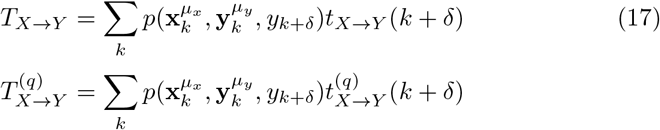

That is, *T*_*X* → *Y*_ and 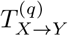 express the average information flow of the local transfer entropies (either using Shannon or Tsallis formulation) with the same probability distribution 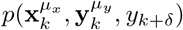.

Higher-order Tsallis local transfer entropies can also be defined:

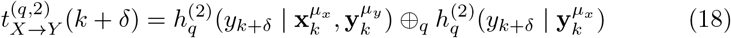

where [39]

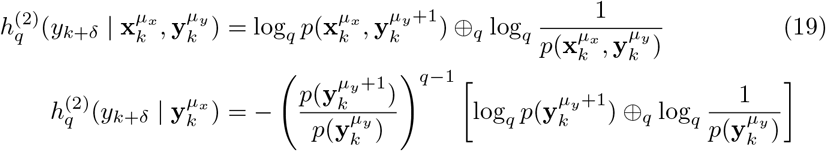

Here, the following relations are used:

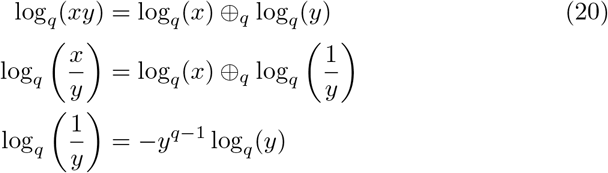

The advantages of using the (*α, q*)-framework are discussed in detail in Supporting Information where two coupled systems are introduced. The first describes the XOR Bernoulli-like noisy channel, which is the case of the non-linearly coupled stochastic random processes, and the second one describes the Gaussian-like noisy channel, which is the example linearly coupled stochastic random processes.

Following Refs. [40, 41, 13], it can be shown that for processes close to the equilibrium, both reversible or irreversible, at constant temperature (see also [42]):

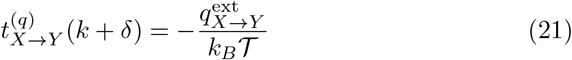

where 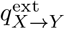 is the heat flow between the system *Y* and the surrounding in the context of the source *X*. The sign minus indicates the heat flow into the physical system *Y* attributed to the surrounding *X* and the local transfer entropy is the opposite. Eq. 21 gives the minimum amount of energy that a physical system (characterised by the dynamical component *Y*) must dissipate into the environment (characterised by the dynamical component *X*) to gain one bit of information. If 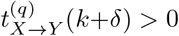, and hence the physical system *Y* gains information, then the heat flows to the surrounding *X* from the system 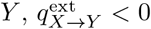. In that case, a measure of the source *X* improves our predictions about the macroscopic transition state of *Y*. For thermally isolated transitions, that is 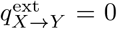, then 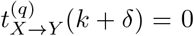, indicating that *Y* and *X* are independent, in other words, *X* and *Y* do not interact with each other.

This study concerns about the random walks of the heat flow in a graph [20], where the primary aim is the direction of the heat flow between any two vertices *X* and *Y*. For that, the following relationship between the local information flow direction and the heat flow direction is defined

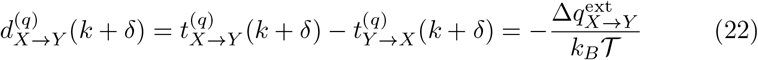

If 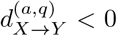, then 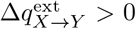 and thus heat flow direction is from *source X* to *target Y*. In Eq. 21 and Eq. 22, 𝒯 is the temperature (in Kelvin) and *k*_B_ is the Boltzmann’s constant.

## 3. Molecular Dynamics Simulations of Modelled Systems

The initial protein structure for molecular dynamics simulations is the BPTI protein with PDB ID 5PTI. It contains 58 amino acids and its structure is investigated by joint X-ray and neutron experiments at a resolution of 1.00 Å [28]. The structure contains three disulphide bonds, namely Cys5-Cys55, Cys14-Cys38, and Cys30-Cys51, as depicted in Fig. 2 and visualised using Visual Molecular Dynamics (VMD) program [43].

**Figure 1:**
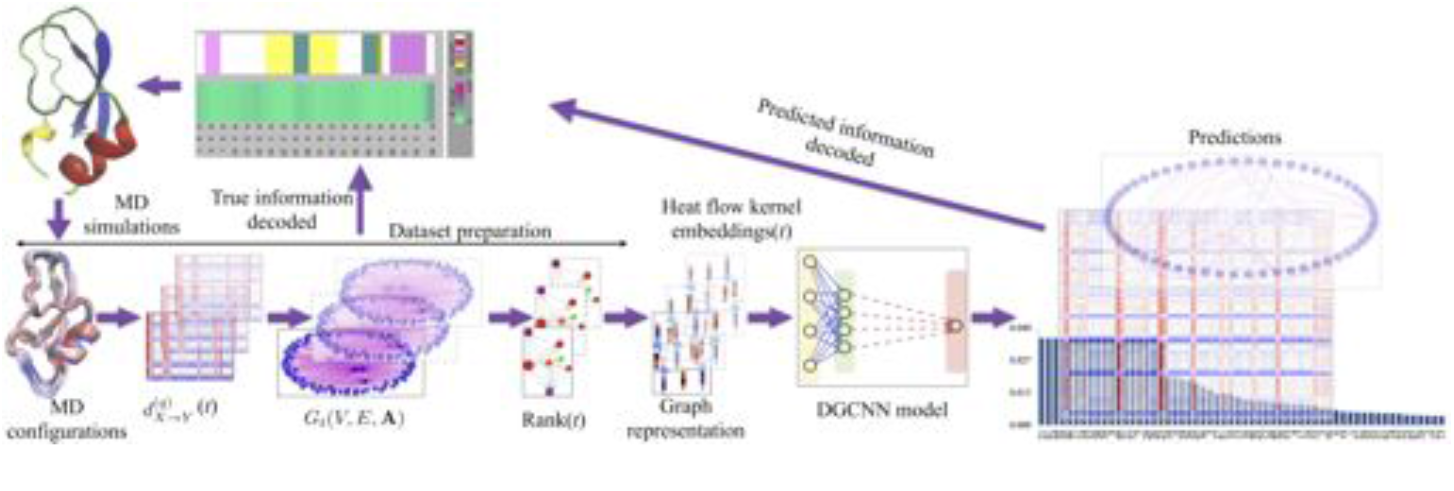
Graphical abstract.

**Figure 2:**
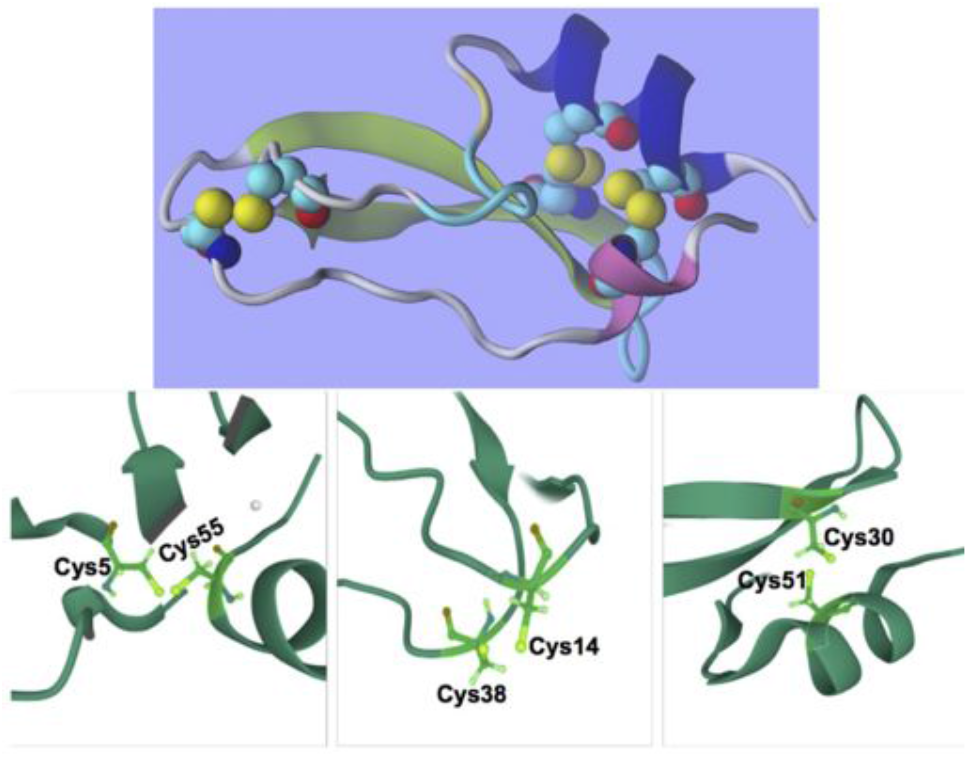
Three-dimensional structure and disulphide bonds of BPTI protein, visualised using VMD program [43].

Next, a partially folded, namely BPTI[14-38]Abu, mutated-type [29] with PDB ID 1YKT was considered at a resolution of 1.70 Å. BPTI[14-38]Abu has a disulphide bond Cys14-Cys38, and Cys5, Cys30, Cys51, and Cys55 are mutated to isometric *α*-amino-*n*-butyric acid. The X-ray structure misses the amino acids Glu57 and Ala58, which are modelled using a three-dimensional structure alignment with BPTI wild-type using VMD program [43] tools.

CHARMM program [44] using the CHARMM36 force field [45, 46, 47] is used for MD simulations. Figure 3 shows the topology of CHARMM36 force field for isometric *α*-amino-*n*-butyric acid and partial charges calculated using quantum mechanics in the gas phase. Other force field parameters, such as bonds, angle bending, torsions, and van der Waals, were chosen based on the atom type as in CHARMM36 [45, 46, 47].

**Figure 3:**
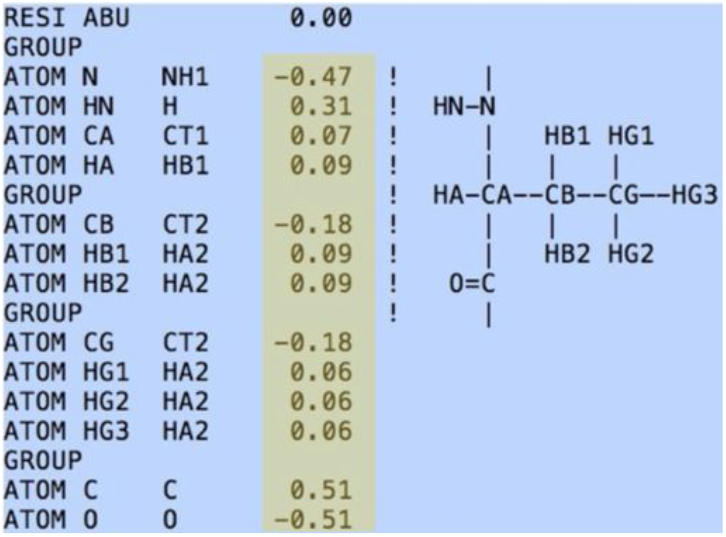
The topology of CHARMM36 force field for isometric *α*-amino-n-butyric acid.

The hydrogen atoms and the protonation state of amino acids were determined using H++ web server ^1^, as in Ref. [48]. TIP3P model [49] water shell is added around the protein with a salt (NaCl) concentration of 0.154 M/l, plus the counter ions to neutralise the systems. Equilibration protocol of the solvent and hydrogen atoms (with heavy atoms of proteins fixed at their X-ray structures) included an energy minimisation (steepest descents and conjugate gradients algorithms [50] for about 5 × 10^3^ − 10 × 10^3^ steps each), heating for about 32 × 10^3^ (*N*; *V*; 𝒯) MD steps (from temperature of 50 K to 800 K in steps of 50 K, constant volume *V* and number of atoms *N*) and cooling for about 40 × 10^3^ (*N*; *V*; 𝒯) MD steps (from 800 K to 300 K in steps of 25 K, constant *V* and *N*). Next, the restraints (*N*; *P*; 𝒯) MD simulations at constant temperature 𝒯 = 300 K and pressure *P* = 1 atm were performed for about 500 ps for each system to equilibrate the solvent and protein by gently lowering the restraints until the density of the solvent converged to the value of 1 gram/cm^3^.

The (*N*; *P*; 𝒯) MD simulations at constant 𝒯 = 300 K; *P* = 1 atm continued for about 2.5 ns for each system without any restraints to finalise the equilibration phase. The production run (*N*; *P*; 𝒯) MD simulations at constant 𝒯 = 300 K; *P* = 1 atm were performed for ten ns for each system. In total, about 30 ns of simulations and around 3 months of CPU time (4 CPUs, 2.8 GHz clock rate) were estimated. A cutoff radius of *R*_cut_ = 12 Åwas used for non-bonded long-range interactions and the SHAKE algorithm [51] to constraint the covalent bonds involving the hydrogen atoms, allowing to use a numerical integration time-step of Δ*t* = 2 fs.

Particle-Mesh-Ewald was employed for long-range electrostatics and the potential shift for handling the short-range van der Waals interactions [42]. (Note that the van der Waals potential and the forces for this choice approach zero as the intra-distances become *R*_cut_, avoiding difficult computations of switching functions and steep forces at large distances, as suggested in Ref. [44]). A fixed dielectric constant of unity was taken and simple potential cutoff approach was used for handling the short-range electrostatic interactions. The Ewald summation was performed with = 4.0/*R*_cut_, as a trade-off between the computation cost and accuracy [42].

The cubic simulation box was used for the periodic boundary conditions, the neighbour list algorithm [52], based on the atom list, with a spherical radius of 13 Å used for keeping the non-bonded list of atoms, and the cutoff radius of the nearest neighbour was 11 ÅHere, a heuristic testing is done every time the energy and forces are calculated and a list update is done if necessary. (Note that this is both safer and more economical than frequency-updating [44]; furthermore, BYCBIM algorithm was employed to update the neighbour lists, as suggested in CHARMM manual [44]). MD simulations configurations of each system were saved every 0.2 ps and for analysis the proteins configurations were saved every 1 ps after removing the overall translational and rotational motions, based on the group of atoms chosen for further computational analysis. Wild-type has around *N* = 19562 atoms and BPTI[14-38]Abu around *N* = 18244 atoms.

For computation of information flow, the fluctuations of coordinates of some property of interest for a group of atoms of each amino acid are stored every picoseconds as a matrix (*T* × 3*n*_*i*_) of time series:

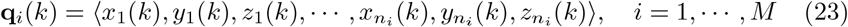

where *n*_i_ is the number of the group of atoms selected from the amino acid *i* and *M* is the total number of amino acids in the protein sequence. Here, *k* is a time index, *k* = 0, 1, ⋯, *T* − 1 where *T* is the number of saved configurations. Then, we tested different properties of the amino acids, such as dipole fluctuations of the side chains, centre of the mass of amino acids, principal component analysis (PCA), and independent component analysis (ICA) to reduce the dimensionality of the vector **q**_i_:

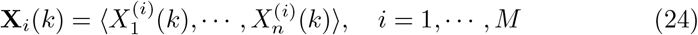

where, typically, 1 ≤ *n* ≤ 3 and *k* = 0, 1, ⋯,*T* − 1. For the PCA and ICA computations, we used the Python software’s classes *PCA* and *FastICA* of the package *sklearn.decomposition*. In our results presented in the following discussion, we used ICA projected coordinates with *n* = 3 using only the backbone atoms Cartesian coordinates of each amino acid. Then, the time delayed embedding method was used to reconstruct the dynamics of the random time processes described by the original vectors **q**_i_(*k*), as in Ref. [20] (and the references therein), *μ*_i_ = (*m*_i_, *τ*_i_). For instance, for any two amino acids *i* and *j*, we construct the vectors 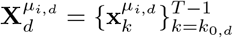, where

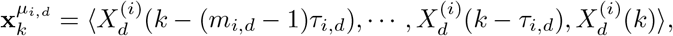

and 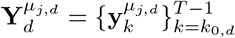:

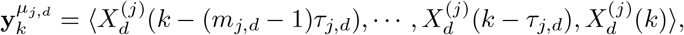

respectively for 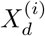 of *i* and 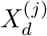 of *j* (for every *d* = 1, 2, ⋯, *n*). Here, *k*_0,d_ = max{(*m*_i,d_ − 1) *τ*_i,d_, (*m*_j,d_ − 1) *τ*_j,d_} to create vectors of the same length.

## 4. Results and Discussion

### 4.1. Structure Stability

Figure 4 shows the root mean square fluctuation (rmsf) of an amino acid in the wild-type BPTI and [14-38]Abu with Ala11Thr, Arg15Lys, Lys26Pro, and Ala27Asp mutations, based on C_*α*_ atom positions. The residues indexed between 1 and 8 show higher fluctuations in BPTI[14-38]Abu than the BPTI-WT structure, indicating a decrease in stability of the 3_10_-helix secondary structure. Similarly, the *α*-helix (residues indexed between 50 and 57) becomes unstable in the structure of BPTI[14-38]Abu (as seen in Fig. 4). Besides, the root mean square fluctuation analyses indicate a decrease in stability of the residues 24-29 of the turn-like secondary structure linking the two *beta* sheets. Interestingly, the coil residues indexed 11 to 18 have fewer fluctuations in BPTI[14-38]Abu than in wild-type BPTI, indicating an increase in the stability of the first *β*-sheet secondary structure. The same increase in stability is observed for turn structure between residues 38 and 42 in mutated BPTI[14-38]Abu protein.

**Figure 4:**
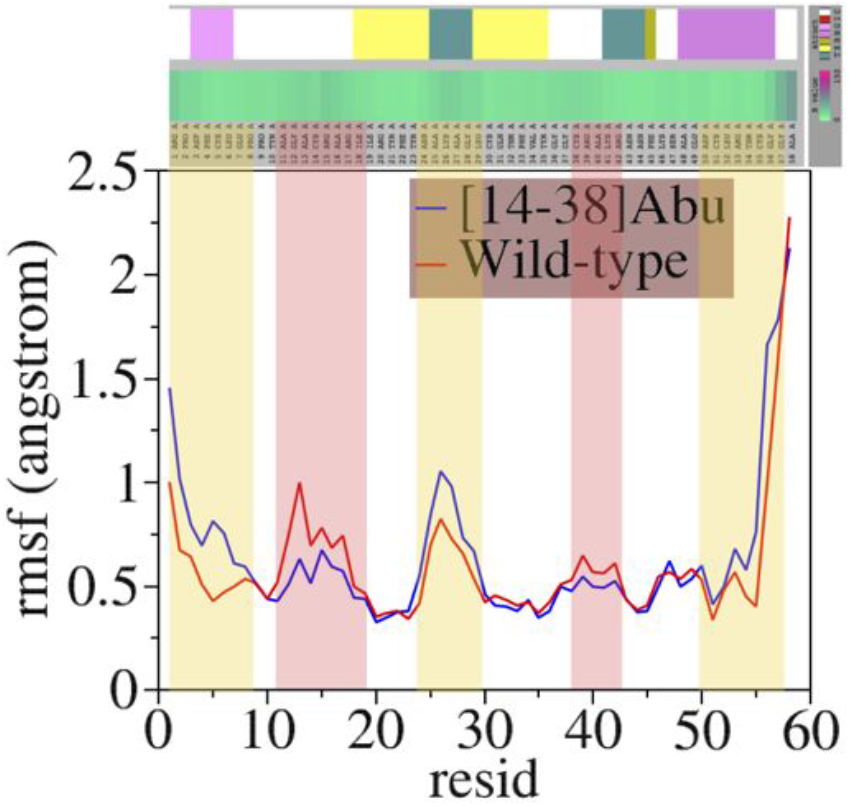
The rmsf of the wild-type BPTI and [14-38]Abu with Ala11Thr, Arg15Lys, Lys26Pro, and Ala27Asp mutations.

Next, the nuclear Overhauser enhancement (the so-called NOE) was computed to determine the spin-lattice dipolar relaxation contributions [53], using CHARMM program [44] implementation. Figure 5 presents the spin-spin NH—HN sequential amino acid NOEs for wild-type BPTI (bottom) and mutated BPTI[14-38]Abu (top). The most interesting is the role of residues Ala16 and Ala58. That is, in the [14-38]Abu protein NOE of Ala16 is positive, and its NOE in wild-type protein is negative; in contrast, in the [14-38]Abu protein NOE of Ala58 is negative, and it changes to a positive value in the wild-type protein. That is due to the increase of the intense of medium-range NOE of Ala16 in BPTI[14-38]Abu protein, which was confirmed by computing NH—NH non-sequential Ala16 NOEs with other amino acids; from our results, Ala16N—Cys38N proton pair was the most intense NOE of 1.077, which are two residues of the coil region linking two β-sheets. This proton pair’s so-called effective relaxation time was 236.89 ps in BPTI[14-38]Abu protein. Note that distance between the pair Ala16N—Cys38N from the X-ray structure is 6.83 Å [29] in mutated [14-38]Abu.

**Figure 5:**
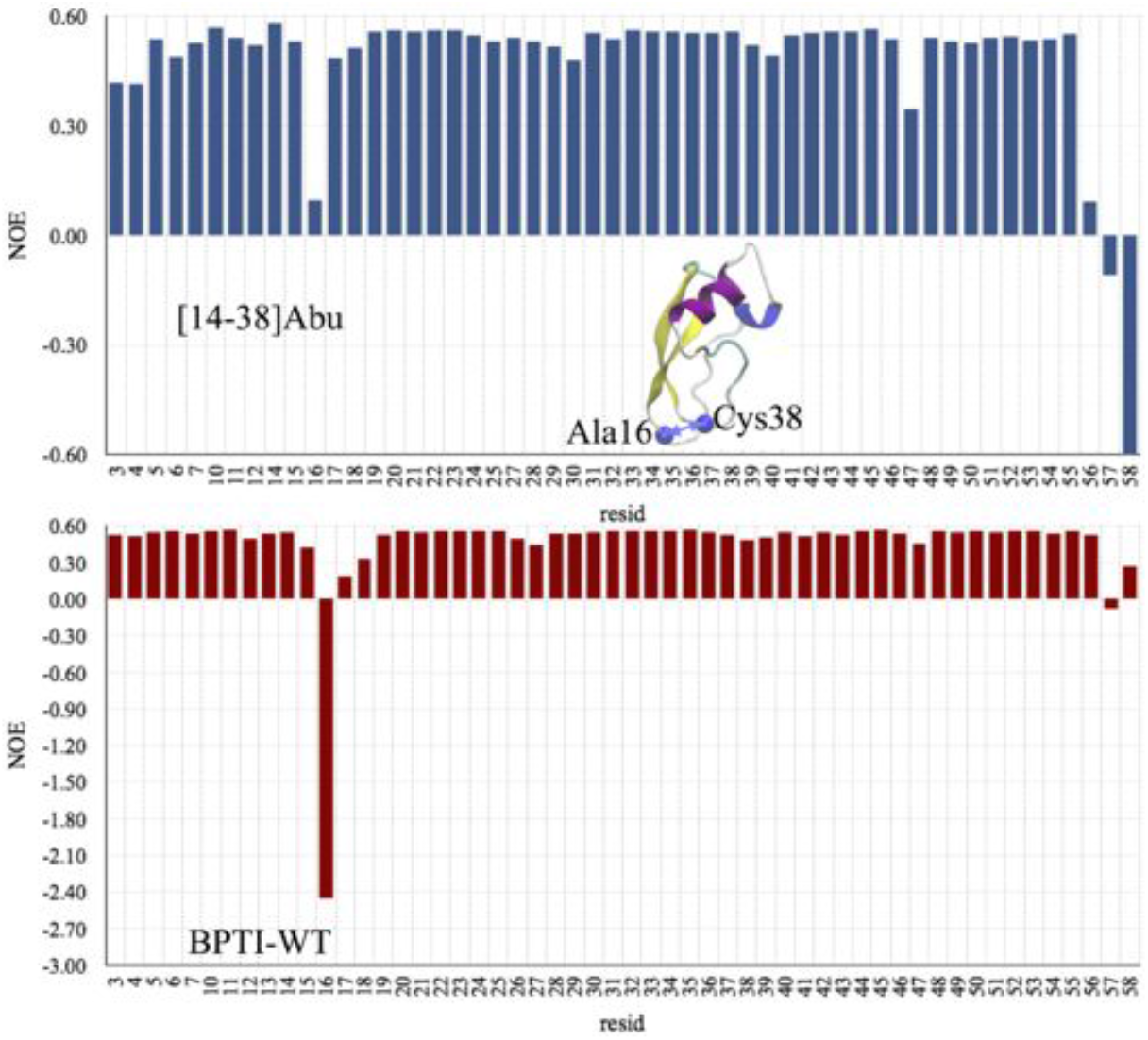
The NOE of the wild-type BPTI (Bottom) and [14-38]Abu with Ala11Thr, Arg15Lys, Lys26Pro, and Ala27Asp mutations (Top).

Furthermore, Table 1 shows the NMR spectra, according to formulas in reference [54]. The CHARMM program [44] was used to compute the turn secondary structure residues 25-28 using the medium-range proton pairs N—N. The equilibrium distances *d* are according to crystal structure experiments from references [28, 29]. Our results indicate that the NOE are more vital for the wild-type BPTI than mutated BPTI[14-38]Abu. These results support experimental findings [55] that residues 25-28 stabilise turn conformations that nucleate folding, which may prevent misfolding of *β*_1_ and *β*_2_ sheets.

**Table 1:** The NOE, effective relaxation time *τ*_*e*_, and TMXE (which is the first time when the time correlation C(t) is less or equal to order parameter of the spin-spin bond), according to formulas in reference [54] and computed using CHARMM program [44], for the turn secondary structure residues 25-28 using the medium-range proton pairs N—N. The equilibrium distances d are according to crystal structure experiments from references [28, 29].

| Proton pair | NOE | $\tau_e$ [ps] | TMXE [ps] | $d$ [Å] |
| --- | --- | --- | --- | --- |
| BPTI-WT |  |  |  |  |
| Ala25N-Lys26N | 1.079 | 222.00 | 2094.0 | 2.72 |
| Ala25N-Ala27N | 1.078 | 136.00 | 1546.0 | 4.38 |
| Ala25N-Gly28N | 1.077 | 12.73 | 194.0 | 4.69 |
| Lys26N-Ala27N | 1.077 | 9.27 | 159.0 | 2.83 |
| Lys26N-Gly28N | 1.075 | 12.00 | 154.0 | 4.24 |
| Ala27N-Gly28N | 1.076 | 10.89 | 163.0 | 2.70 |
| BPTI[14-38]Abu |  |  |  |  |
| Ala25N-Lys26N | 1.075 | 24.49 | 211.0 | 3.04 |
| Ala25N-Ala27N | 1.071 | 36.29 | 279.0 | 4.77 |
| Ala25N-Gly28N | 1.075 | 44.59 | 439.0 | 4.55 |
| Lys26N-Ala27N | 1.074 | 42.33 | 376.0 | 3.01 |
| Lys26N-Gly28N | 1.077 | 51.49 | 474.0 | 3.99 |
| Ala27N-Gly28N | 1.076 | 39.12 | 280.0 | 2.62 |

Moreover, Table 2 presents the NMR spectra for the 3_10_-helix secondary structure residues 3-6 using the medium-range proton pairs N—N. NHs cross peaks are sharper in the mutated [14-38]Abu protein than wild-type BPTI, indicating that these proton pairs are not in an intermediate exchange time regime and do not sample multiple conformations. These findings agree with NMR experimental results [55, 56].

**Table 2:** The NOE, effective relaxation time *τ*_*e*_, and TMXE, according to formulas in reference [54] and computed using CHARMM program [44], for the 3_10_-helix secondary structure residues 3-6 using the medium-range proton pairs N—N. The equilibrium distances d are according to crystal structure experiments from references [28, 29].

| Proton pair | NOE | $\tau_e$ [ps] | TMXE [ps] | $d$ [Å] |
| --- | --- | --- | --- | --- |
| BPTI-WT |  |  |  |  |
| Asp3N-Phe4N | 1.079 | 297.5 | 2162.0 | 2.83 |
| Asp3N-Cys5N | 1.078 | 109.0 | 1068.0 | 4.41 |
| Asp3N-Leu6N | 1.075 | 83.4 | 639.0 | 5.60 |
| Phe4N-Cys5N | 1.078 | 238.0 | 2246.0 | 2.72 |
| Phe4N-Leu6N | 1.077 | 132.9 | 1687.0 | 4.38 |
| Cys5-Leu6N | 1.076 | 55.2 | 949.0 | 2.71 |
| BPTI[14-38]Abu |  |  |  |  |
| Asp3N-Phe4N | 1.086 | 233.2 | 1032.0 | 2.85 |
| Asp3N-Abu5N | 1.082 | 182.5 | 996.0 | 4.29 |
| Asp3N-Leu6N | 1.072 | 83.6 | 304.0 | 5.42 |
| Phe4N-Abu5N | 1.084 | 351.2 | 2049.0 | 2.71 |
| Phe4N-Leu6N | 1.074 | 79.3 | 347.0 | 4.34 |
| Abu5-Leu6N | 1.072 | 71.6 | 349.0 | 2.67 |

### 4.2. Free Energy Change Upon Mutations

Figure 6 presents the thermodynamic cycle for calculation of binding free energy of an amino acid in a protein relative to a reference surrounding phase [57]. (Note that the same thermodynamic cycle is used in the previous studies [58, 59] using an in-house code; however, in this study, the CHARMM program [44] is used). The differences in the solvation free energy (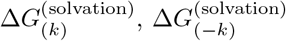, and 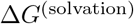) in reference surrounding phase (often gas phase) and in a solvent (such as salted water) are computed using Poisson-Boltzmann (PB) method [42] implemented in CHARMM [44]. Here, 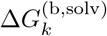 and 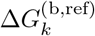 denotes the binding free energies of the amino acid *k* in the solvent and gas phase, respectively. 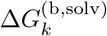 is computed as:

**Figure 6:**
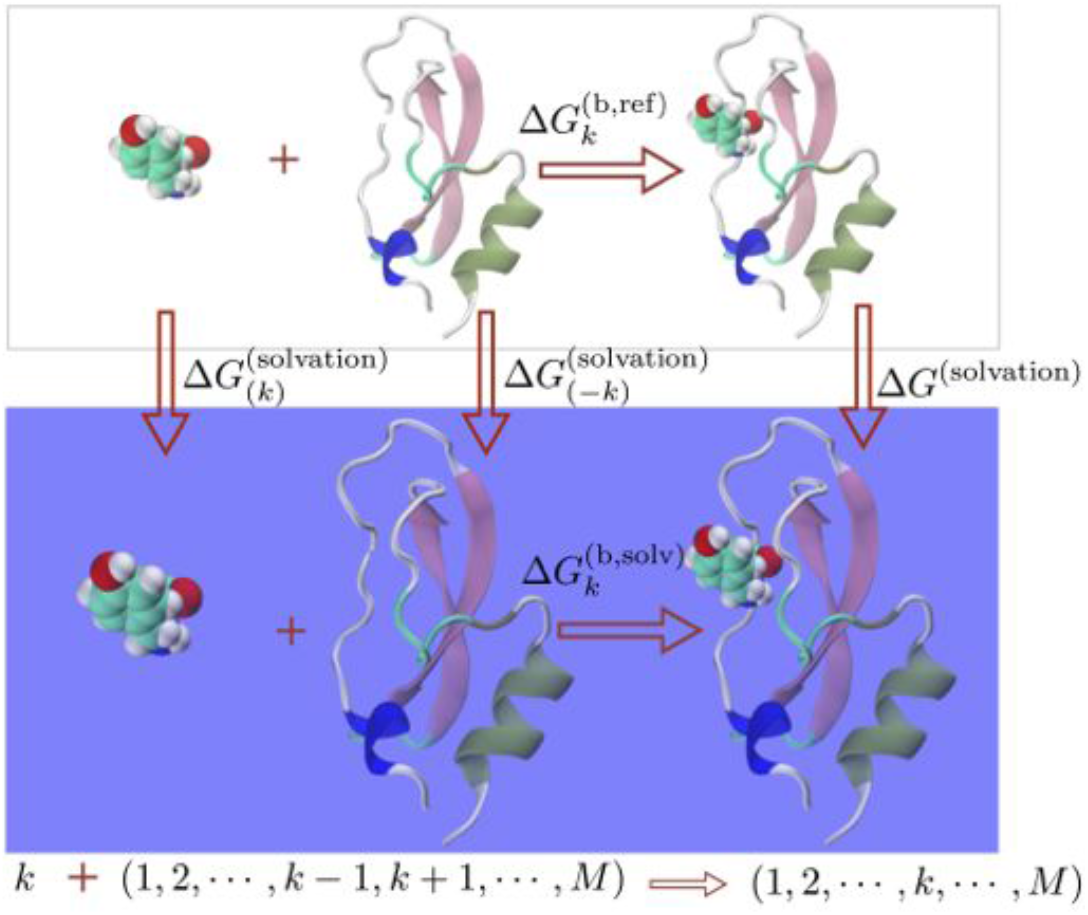
Thermodynamic cycle for calculation of binding free energy of an amino acid in a protein. The differences in the solvation free energy, 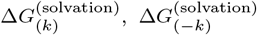, and 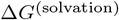 in reference surrounding phase (often gas phase) and in solvent (such as water and salt).

**Figure 7:**
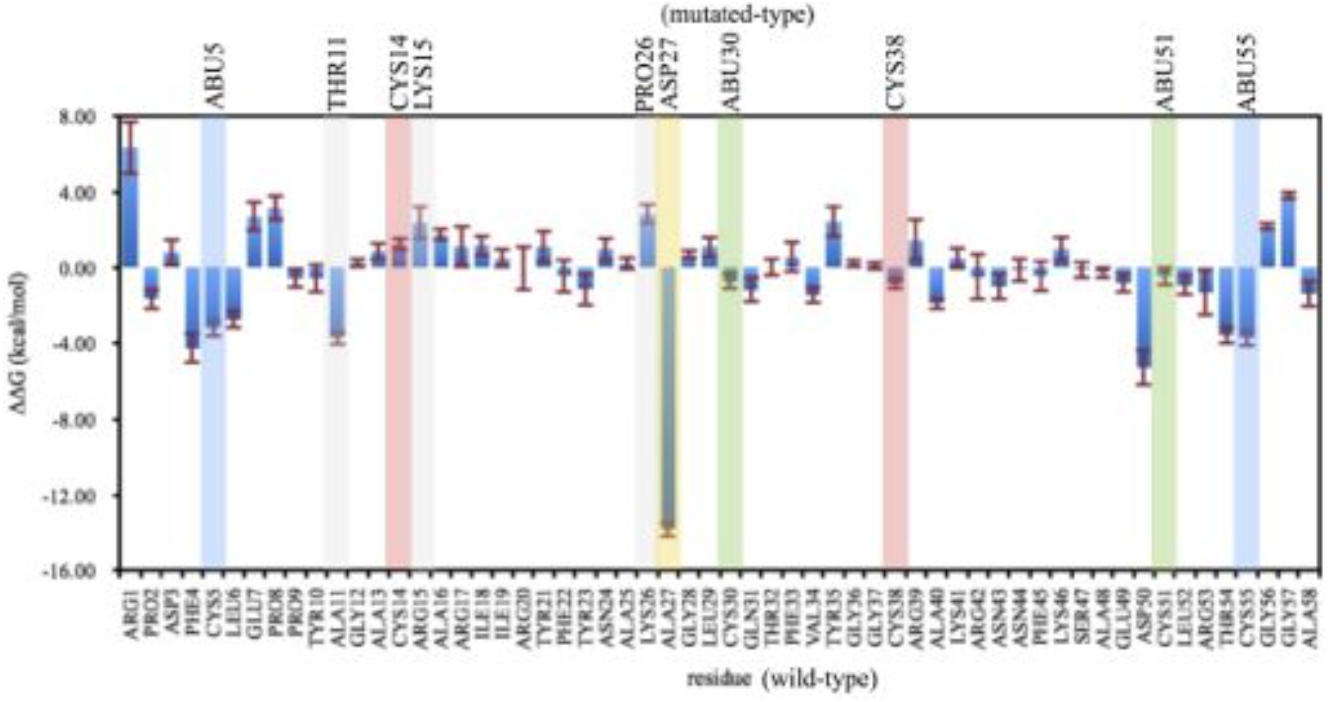
The free energy changes for each amino acid upon mutation from protein BPTI-WT to BPTI[14-38]Abu.

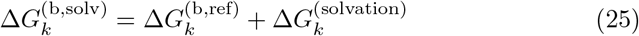

where

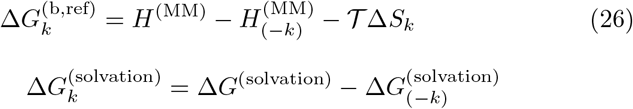

Here, *H*^(MM)^ and 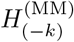 represent the potential interaction energy of the entire system and subsystem without amino acid *k*, based on the CHARMM36 molecular force field. Furthermore, Δ*S*_k_ is the change on the entropy with 𝒯 being the temperature (typically, 𝒯 = 300 K, the same as in MD simulations), which is decomposed into three terms, namely rotational, translational and configurational entropies, implemented in CHARMM [44]. Note that, here, Andricioaei & Karplus formula is used to compute configurational entropy [60, 61].

Moreover, 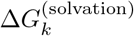 is decomposed into the polar (or electrostatic) and non-polar (or solvent-accessible-surface area, SASA) terms as:

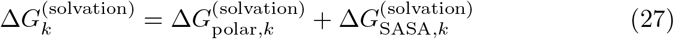

In Eq. 27, 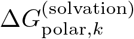 is calculated using PB method and 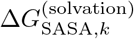 is parametrised as

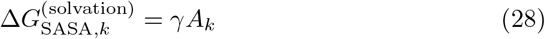

where *A*_k_ is the value of SASA for the amino acid *k* present in the protein and γ= 0.006 kcal × mol^−1^ × Å−2. The coefficient γand parametrisation of the surface effects are designed such to include short-range effects, dominated by the first solvation shell [62, 63]. There is a wide range of the values of γbecause of different models used to represent the molecular surface [62]. However, small values of γindicate that molecular surface contribution has only a small impact on the free energy changes due to large-scale conformation changes. Note that all the computed quantities are time averaged ones over different configurations obtained from MD simulations; typically, 2000 configurations were used. The statistical analyses use an 95 % confidence interval for the mean [64]. All the CHARMM’s scripts for these statistical analyses and computations are specifically created in this study with the help of CHARMM [44] manual and Forum website ^2^ over several years.

The computations included each system (such as BPTI-WT and BPTI[14-38]Abu proteins). The differences between the free energy changes due to mutations are calculated as:

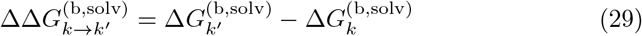

where *k* is the amino acid in BPTI-WT and *k*^’^ is the amino acid with the same ID in BPTI[14-38]Abu.

Figure 8 shows the results for 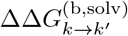. In the computations of the electrostatic Poisson-Boltzmann terms, the dielectric constants of the protein and water are, respectively, ϵ_p_ = 2 and ϵ_w_ = 80. The solvent salt concentration was 0.154 M/l (the same as in atomistic MD simulations). From our calculations, mutation of Cys5 and Cys55 to Abu resulted in favour of the stability of the structure, respectively, 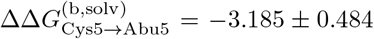 kcal/mol and 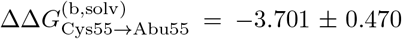 kcal/mol. On the other hand, the mutations of Cys30 and Cys51 to Abu favoured much less the stability of the BPTI[14-38]Abu; in particular, 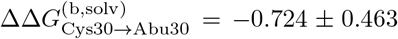 kcal/mol and 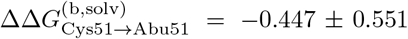 kcal/mol. Interestingly, the most favoured mutation in enhancing the stability of BPTI[14-38]Abu structure is Ala27⟶Asp27 with a value of 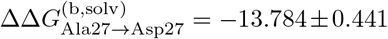 kcal/mol. In contrast, the mutation of Lys26 to Pro26 disfavoured the stability with a value of 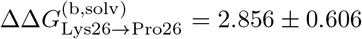 kcal/mol; similarly the mutation of Arg15 to Lys15 with a value of 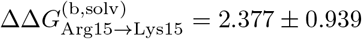 kcal/mol.

**Figure 8:**
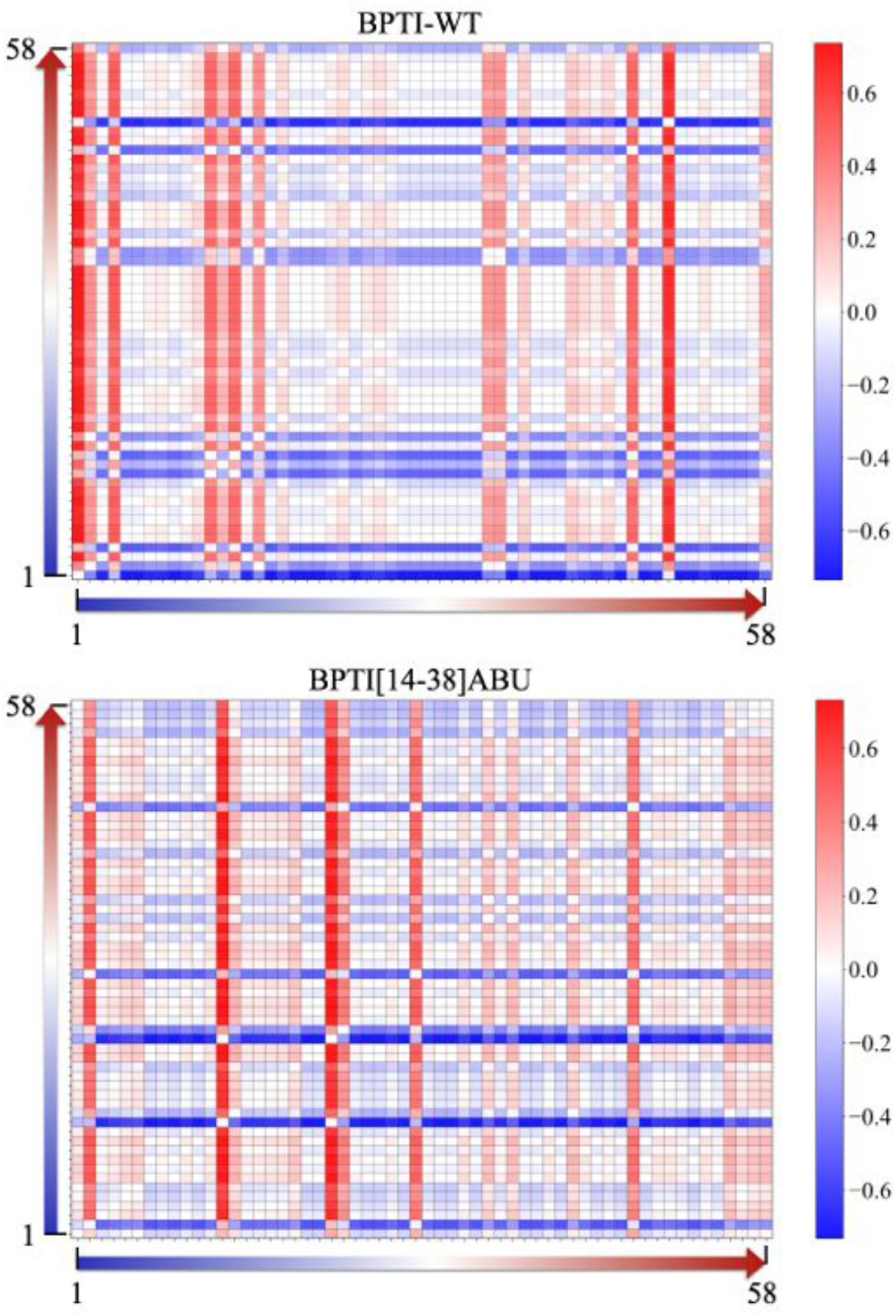
The directional information flow (in nats) for BPTI wild-type (Top) and BPTI[14-38]Abu (Bottom).

Furthermore, the mutation of Ala11 to Thr11 favoured the stability of BPTI[14-38]Abu structure with a change on the free energy of −3.708 ± 0.400 kcal/mol. Interestingly, other amino acids, which are not involved in the mutation, contribute to the stability of the protein structure. This indicates that the interaction network topology between the amino acids in the protein sequence has possibly changed upon the mutations. Therefore, the following graph information flow network analyses could potentially enable identification of the changes in the amino acids interaction networks of both proteins. The aim is to reveal the encoded information of the sequence of amino acids into a protein structure using the newly developed (*α, q*) information-theoretic measures. In particular, the primary interest is to investigate statistical efficiency of the Sharma-Mittal transfer entropy, based on the (*α, q*)-framework.

### 4.3. The (a, q)-Framework Directional Information Flow

In this study, the information flow direction is calculated from the Sharma-Mittal symbolic encoding transfer entropies, based on the (*a, q*)-framework, as:

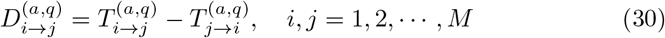

where *M* is the number of amino acids in the structure. The choice was *α* = 2.0 and *q* = 0.85, based on the analysis presented in Supporting Information material associated with study. Here, if 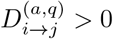 then information flows from source *i* to target *j*, or *i* influences *j*. Figure 8 presents heat map plots of 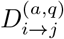 (in nats) for BPTI-WT and BPTI[14-38]Abu, where a red colour indicates a strong *source* of information and blue one indicates a *target* (or *sink*) amino acid of information flow. Our results clearly identify the source and target residues and the change in the amino acid’s propensity to be a *source* or *target* (*sink*) of information flow for BPTI-WT and BPTI[14-38]Abu.

Figure 9 shows the differences in information flow for each amino acid in BPTI wild-type and BPTI[14-38]Abu structures (Fig. 9(a)) calculated as:

**Figure 9:**
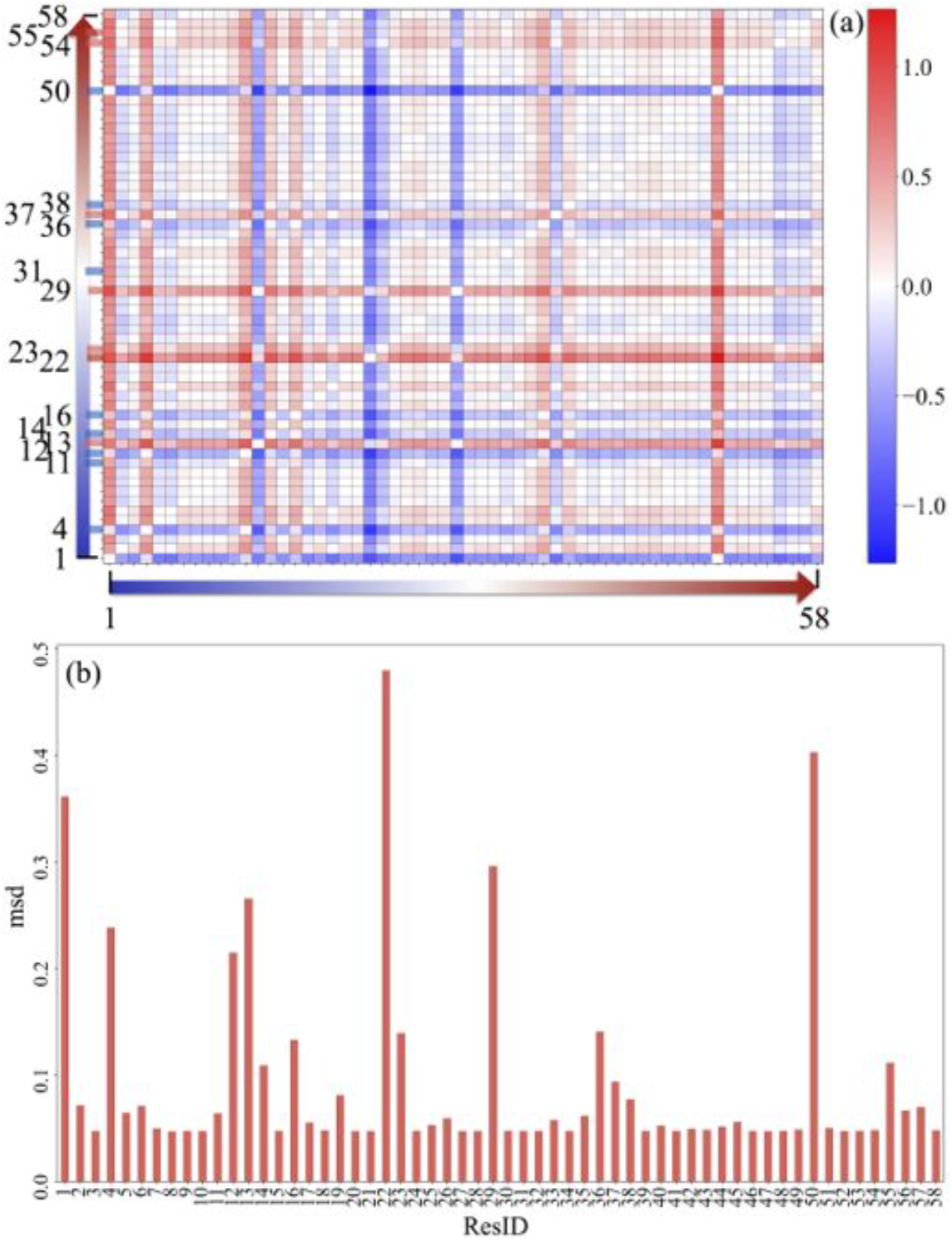
(a) The directional information flow difference between BPTI wild-type and BPTI[14-38]Abu; (b) The msd of the information flow difference between BPTI wild-type and BPTI[14-38]Abu.

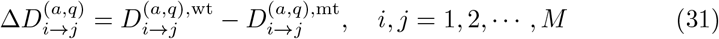

Here, if 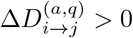, then amino acid *i* has a greater propensity to be a source of information than *j* in wild-type structure relative mutated protein, which is indicated in Fig. 9(a) by a red colour; otherwise, if 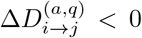, then *i* has a greater propensity to be a target of information than *j* in wild-type protein relative to mutated one, indicated in Fig. 9(a) by a blue colour. In Fig. 9, we have depicted the amino acids with the greatest positive impact as a *source* of information flow, such as Ala13, Phe22, Tyr23, Leu29, Gly37, Thr54, and Cys55(wild-type)/Abu55(mutated-type). Furthermore, the amino acids with the greatest positive impact as a *sink* of information flow include: Arg1, Phe4, Ala11(wild-type)/Thr11(mutated-type), Gly12, Cys14, Ala16, Gln31, Gly36, Cys38, and Asp50.

We also calculated the mean-square-deviation (msd) of 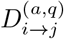 between BPTI-WT and BPTI[14-38]Abu as:

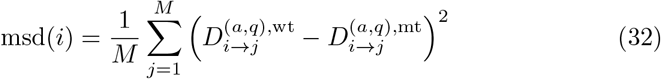

The results are presented graphically in Fig. 9(b), where one can clearly identify the amino acids with the largest deviation in 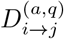 from wild-to mutated-type protein. Interestingly, there is some relationship between the changes on 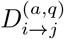 and the changes on the free energies (magnitude); for example, large values of msd for residues 1, 4, 11, 50, and 55 correspond to significant changes on the free energies. That may indicate that the information-theoretic measure 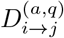 (based on the fluctuations of the encoded information) captures the nature of interactions within a protein structure, which is highly non-linear based on the physics-nature of molecular force fields [3]. This is possibly related to the fact that (*α, q*)-framework maximises the average information flow between the Markovian time processes, as was analysed in the Supporting Information for more intuitive non-linearly coupled random Markovian processes (such as the input-output of the XOR Bernoulli-like noisy channel).

### 4.4. Amino Acids Ranking

Next, we considered graph-theoretic measures (such as PageRank, Authority, and Hub) [65, 66, 67, 68, 69, 70] to rank the impact of the amino acids in the protein’s interaction network; for that, first, a DiGraph is constructed using directional information flow matrix 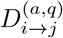. For the PageRank, the algorithm introduced in Ref. [20] is used. For the Authority and Hub ranks, a new algorithm is developed based on the so-called Google matrix, computed as in Ref. [20]. The algorithm uses the Google matrix *L* as an input, then, iteratively, the Authority rank (*a*_i_, for *i* = 1, 2, ⋯, *n*) and Hub rank (*h*_i_, for *i* = 1, 2, ⋯, *n*) scores are computed for each vertex *i* of the DiGraph *G* [68]:

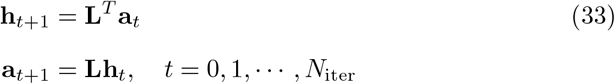

where ()^T^ stands for the transpose, *t* is an iteration index, and *N*_iter_ is the number of iterations (maximum 100 considered in this study). At each iteration, the vectors **a** = (*a*_1_, *a*_2_, ⋯, *a*_n_)^T^ and **h** = (*h*_1_, *h*_2_, ⋯, *h*_n_)^T^ are normalised to one. Here, the Authority score indicates the rank of an amino acid as a *source* of information flow through the graph and the Hub scoring ranks the vertices (and thus the amino acids) based on their influence as *mediators* or *gateways* of the information flow along a path (identified in this study as the *shortest* and the *best* paths, as described below) from a *source* to the corresponding *target*.

Our results of the PageRank and Hub ranks for each amino acid of BPTI wild-type and BPTI[14-38]Abu are shown in Fig. 10. The PageRank and Hub scores may show a helpful trend with the changes in the free energy from the wild-to the mutated type. In particular, an increase in the PageRank’s rank of an amino acid, relative to others, from BPTI-WT to BPTI[14-38]Abu results in a favourable free energy change of that amino acid in mutated structure. For instance, mutation of Ala27 to Asp27 increases the rank of Asp27 (and so its influence) in BPTI[14-38]Abu, and a favourable free energy change is also observed. Similar results are obtained for amino acids 4, 11, 14, 15, 26, 38, and 50, as indicated in Fig. 10. Interestingly, the same trend is observed when the Hub rank increases; in particular, the increase of the Hub rank scores for the amino acids with ID 4, 5, 6, 30, 54, and 55, relative to others in the same protein structure from the wild-to mutated-type protein, correlates well with the corresponding changes on the free energy. That may have a significant impact on predicting the amino acids that, if mutated, could result in a favourable free energy change due to their cooperative contribution within the structure as a mediator of the directional information flow in the protein interaction network of amino acids.

**Figure 10:**
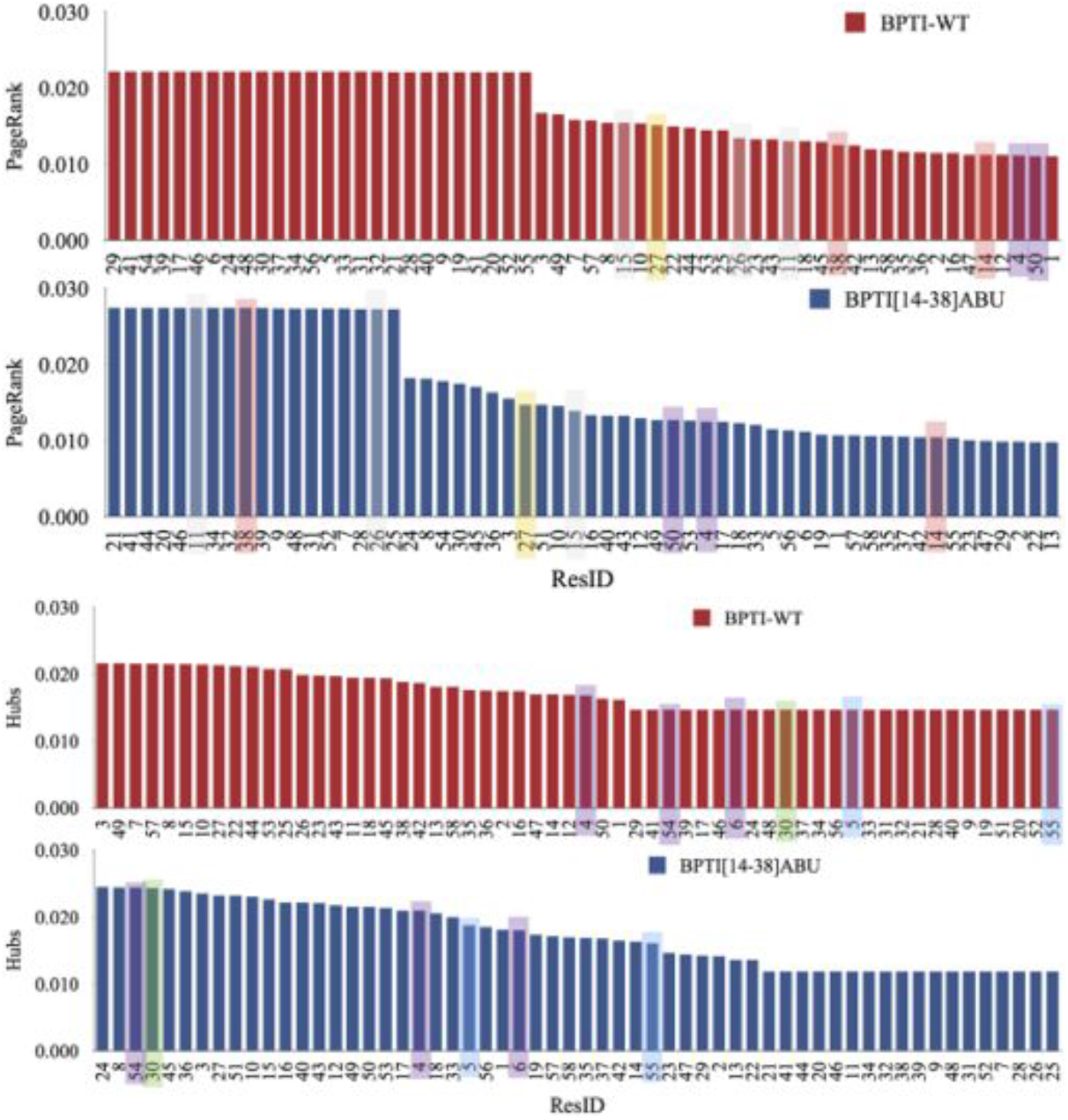
The PageRank and Hub ranking for each amino acid of BPTI wild-type and BPTI[14-38]Abu.

Authority rank of amino acids showed strong correlation with PageRank, as presented in Fig. 11. Therefore, our study indicates that these two DiGraph interaction network measures characterise similar topological properties and their can be used interchangeably.

**Figure 11:**
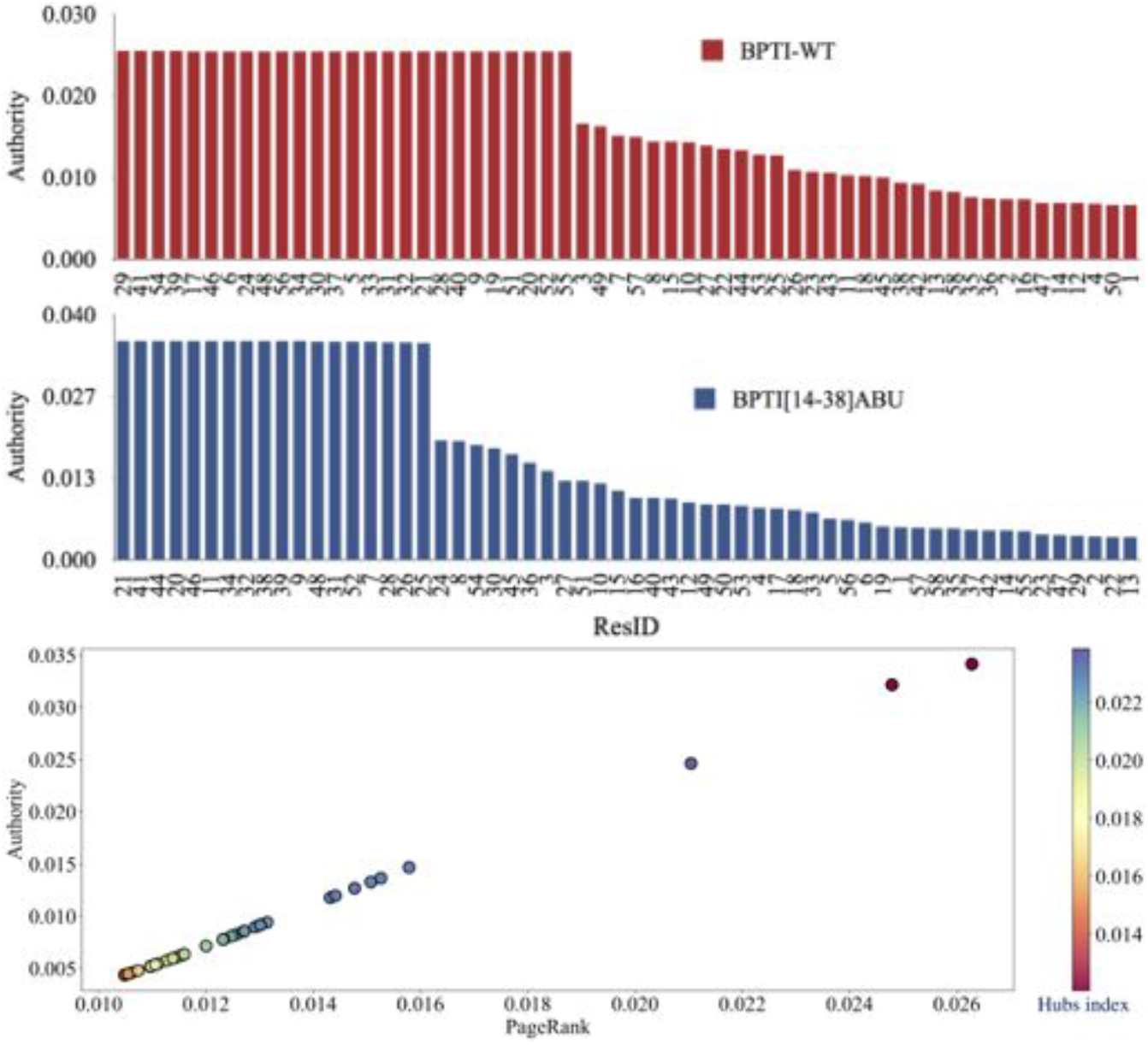
The authority ranking for each amino acid of BPTI wild-type and BPTI[14-38]Abu (two plots in the top) and the correlation between the PageRank and Authority indices (the plot in the bottom).

### 4.5. Amino Acids Information Flow Pathways

The above graph-theoretic measures can also identify the shortest and best paths between a *source* and *target* amino acid.s Here, the algorithm, developed in Ref. [34], is used to determine the shortest and the best paths of information flow in an amino acids’ interaction DiGraph network topology. In Fig. 12, we present the shortest/best paths of amino acids between Cys14 and Cys38 for BPTI wild-type and BPTI[14-38]Abu. The algorithm given in Ref. [71] was used, as implemented in Ref. [34]. The paths are identified by entering the first and the last amino acids, representing, respectively, the so-called above the *target* and *source* of information flow; practically, this is determined by comparing the PageRank ranks, PageRank(first) *<* PageRank(last), or information flows from the last to the first. Figure 11 shows different possible paths between a target and source of the amino acids for BPTI-WT and BPTI[14-38]Abu. All the intermediate amino acids act as hubs (mediators or gateways of the directional information flow from a source to the corresponding target amino acid). It is worth mentioning that all paths for wild- and mutated-type proteins pass through the same number of hubs; however, the mediator role of the Hubs is undertaken by different amino acids. Again, this indicates the changes occurring in the interaction network DiGraph and its directional information flow topology due to the mutation of amino acids.

**Figure 12:**
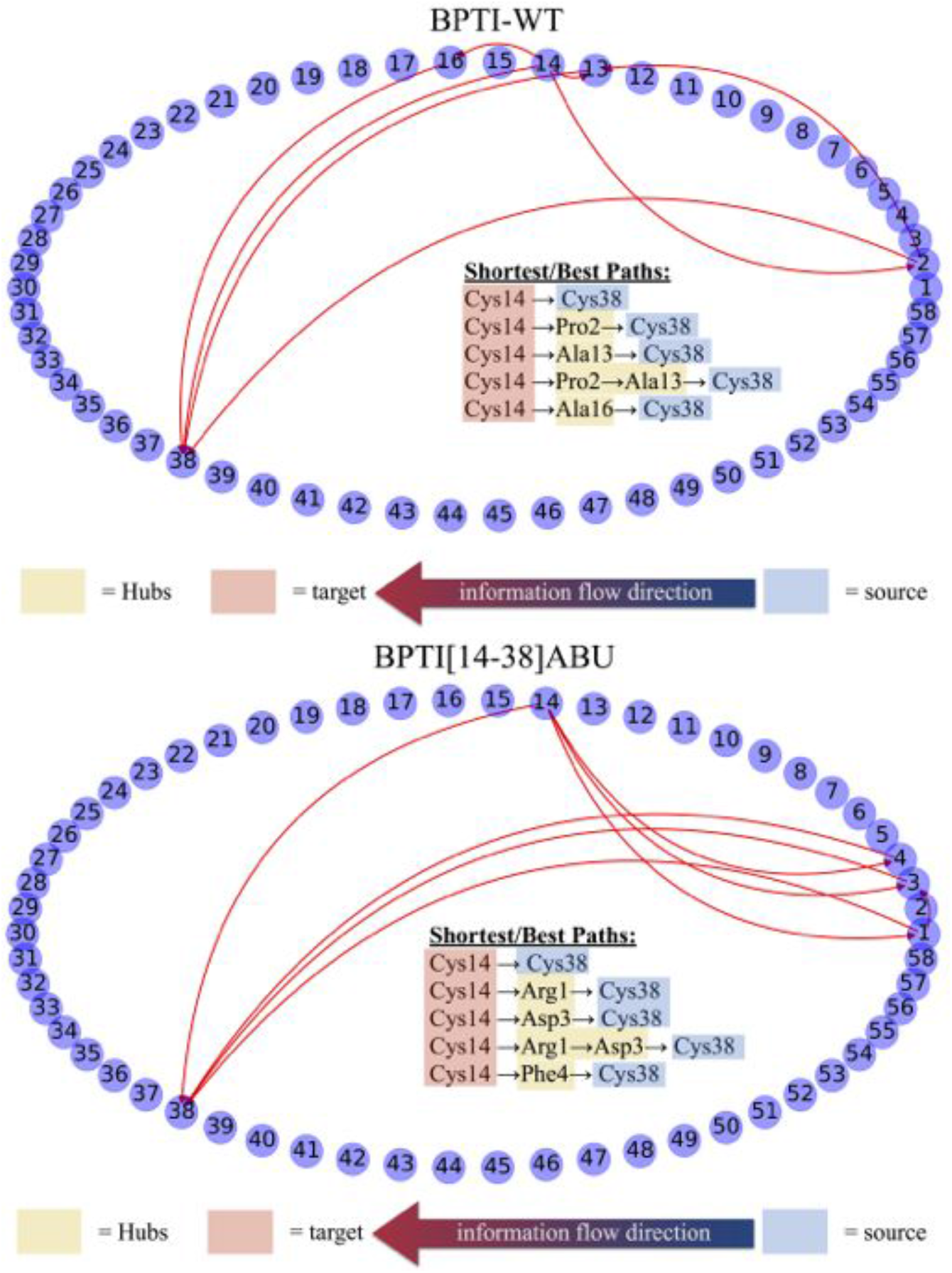
The shortest/best paths of amino acids between residues Cys14 and Cys38 for BPTI wild-type and BPTI[14-38]Abu. The algorithm given in Ref. [71] and implemented in [34] was used.

Figure 13 presents a long path and the corresponding short path of amino acids between Cys14 and Cys38 for BPTI wild-type (see Top plot) and the shortest path between Abu55 and Abu5 for BPTI[14-38]Abu. This kind of examination can be used to identify the most influential Hubs from the source to target in an interaction network DiGraph. Pro2 is potentially the most influential gateway of the information flow between Cys14 and Cys38.

**Figure 13:**
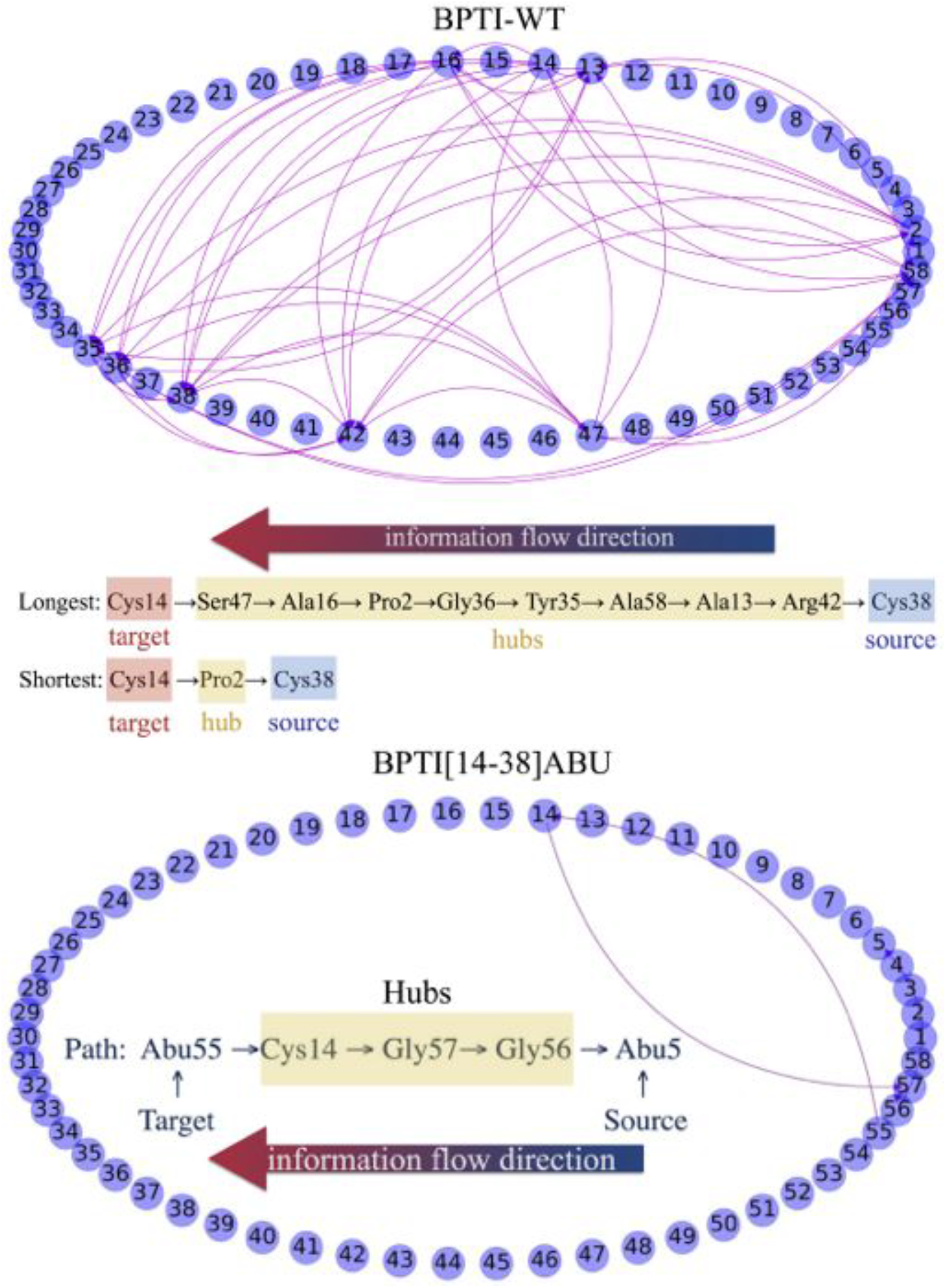
(Top) A long and short path of amino acids between Cys14 and Cys38 for BPTI wild-type; (Bottom) A short path between Abu55 and Abu5 for BPTI[14-38]Abu.

### 4.6. Machine Learning of Amino Acids Ranks

#### 4.6.1. Heat Flow Kernel DiGraph Embedding

Consider a directed graph *G*(*V, E*, **A**) with *M* vertices, *V* = {*v*_1_, *v*_2_, ⋯, *v*_M_}, and *m* edges, *E* = {*e*_1_, *e*_2_, ⋯, *e*_n_} where *e*_i_ = (*u, v*) is a directed edge from *u* to *v*, characterised by the weight *W*_uv_ and an adjacency matrix **A**:

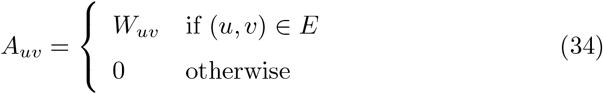

Next, two diagonal matrices can be determined to characterise the in- and out-degrees of vertices, denoted **D**^(in)^ and **D**^(out)^, respectively, as:

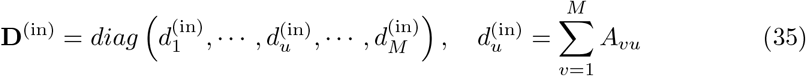

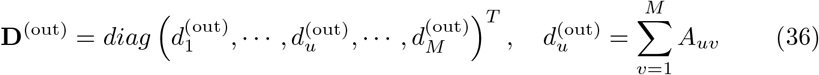

Then, the transition probability matrix 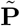of rank *M* × *M* of a random walk on a DiGraph is given as:

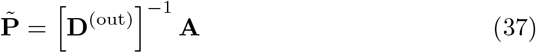

For a strongly connected digraph the random walk characterised by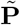 is an irreducible and aperiodic Markovian transition process, which is satisfied if there are no vertices with 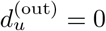 [20]. The following transition probability matrix is suggested to guarantee the irreducibility and aperiodicity

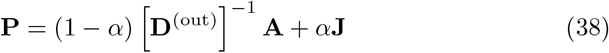

where **J** is a jumping matrix given in terms of the unitary diagonal matrix **1**_M_ of rank *M* × *M* as:

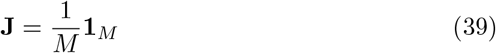

Here, *α* is a number between 0.1 and 0.2 [20] (and the references therein). Then, the normalised Laplacian matrix is constructed as [26]:

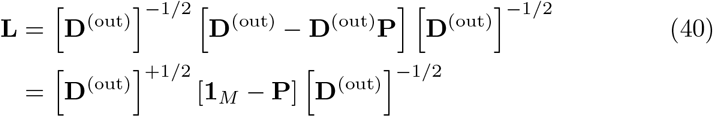

Spectral decomposition of **L** matrix is given as:

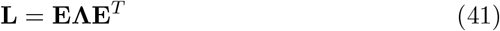

where **E** is the matrix of ordered eigenvectors, 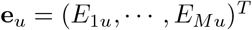 as columns and **Λ** is a diagonal matrix of ordered eigenvalues (*λ*_1_ *< λ*_2_ *<* ⋯*< λ*_M_), which are all non-negative.

The vertices of the DiGraph *G* are mapped into a vector space using the heat kernel such that for each vertex, we determine a so-called embedded vector 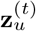 of *M* dimensions at any time *t* (for *u* = 1, 2, ⋯,*M*) [26]:

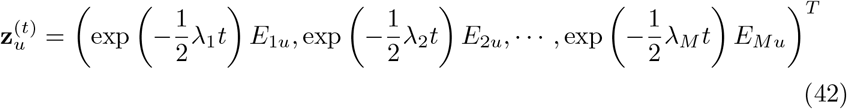

where *t* is the random walk diffusive time of heat flow on the directed graph. That represents a kernel mapping

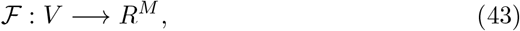

which embeds each vertex of the DiGraph into a vector space *R*^M^. Thus, the Euclidean distance between any two vertices *u* and *v* is defined as:

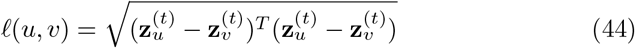

#### 4.6.2. Deep Graph Convolution Neural Network Model

First, we describe the graph representation of the dataset created by computation of the heat flow 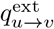 between two amino acids *u* and *v*, representing the vertices of a DiGraph. That is done at regular time intervals of MD simulations; therefore, a set of such graph representations will be stored in the dataset with *t* = 0, 1, ⋯,*T* − 1 entries where *T* is the number of snapshots saved from a typical MD simulation.

Figure 14(a) illustrates a graph representation (adapted from Ref. [25]) of the heat flow at some time frame *t, G*_t_(*V, E*, **A**), in a protein where vertices represent amino acids and the arrow lines indicate the direction of heat flow; that is, if 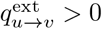, an edge is added from *u* to *v* with weight 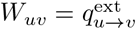. This constructs an adjacency matrix **A**, as illustrated in Fig. 14(b). Each vertex *u* is associated with an embedding vector **z**_u_, constructing the embedding matrix of rank *M* × *M* (see Fig. 14(c)), which is computed from spectral decomposition of the Laplacian **L** matrix using Eq. 42 and **L** is computed using Eq. 40. Besides, one could assign an embedding matrix to the edge attributes; here, the probability transition matrix **P**, given by Eq. 38, is used as the edge embedding matrix, and hence *n* = 1.

**Figure 14:**
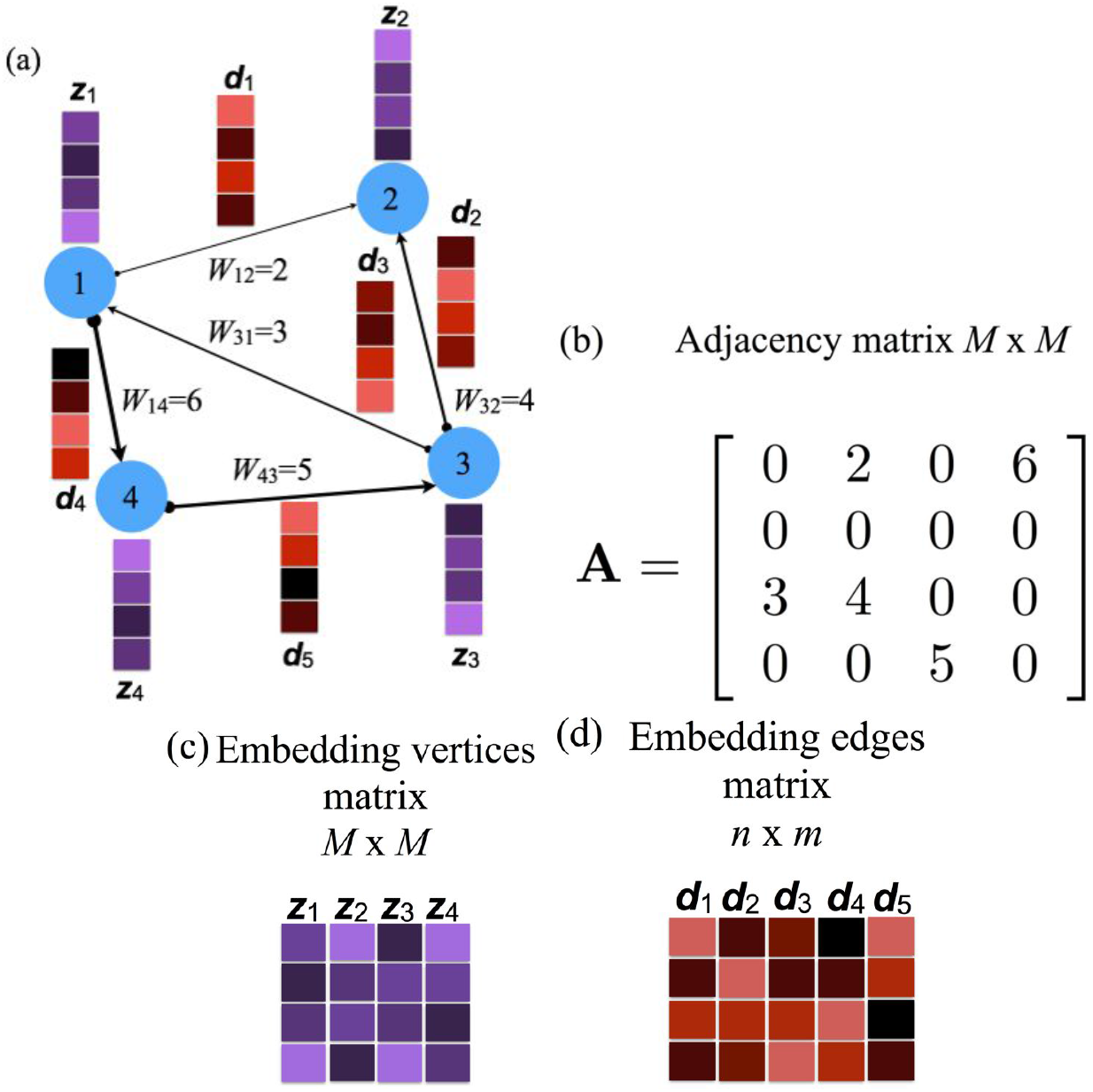
a) Illustration of a graph representation indicating the vertices set, edges set, edges weights, edges embeddings, vertex embeddings; (b) Adjacency matrix *M* × *M*; (c) Embedded vertices matrix *M* × *M*; and (d) Embedded edges matrix *n* × *m*. It is slightly adapted from Ref. [25].

A graph neural network (GNN) is a computation model that uses the graph representation described above as an input (that is, the vertex embedding vectors **z**_u_, *u* = 1, 2, ⋯,*M*, adjacency matrix **A**, and edge embedding matrix **P**) and transfers it into a set of *L* layers. The vertex embedding vectors **z**_u_, at every layer, update to construct the so-called intermediate *hidden* representations **h**_u,l_ with the output vertex embedding vector **h**_u,L_. The GNN uses matrix representation; **Z** is the matrix of vertex embeddings with the columns the vertex embeddings **z**_u_. Similarly, the layer embeddings **H**_l_ and the output matrix **Y** are defined. The input layer of the GNN contains the information about the embeddings of each vertex *u* in **Z** and the last layer (layer *L*) contains the information about the vertices and their context within the graph in **H**_L_.

In this study, we assign to each vertex *u* of the graph an output vector **y**_u_, representing the set of measurements to be predicted using the so-called *regression* at a vertex-level task. In particular, we used the amino acids ranks (such as PageRank, Authority, and Hub ranks):

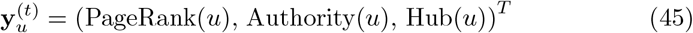

That is done using the graph structure and vertex embeddings. Besides, the loss functions are independently defined at each vertex *u*:

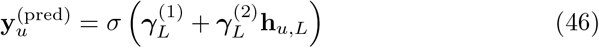

where σ is a sigmoid or softmax function and the column vector 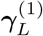 and a matrix 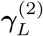 are learned parameters. (Note that a final linear transformation is sometimes needed to map vertex embeddings to the expected output feature dimensionality). The loss function is the mean square error (mse). One can also use the same network to predict the edge attribute embeddings (such as the instant directional information flow 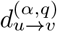) between any two vertices *u* and *v*, such as the transition probability matrix elements [25]:

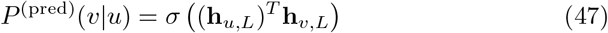

This study uses the graph convolutional network (GCN) among the other GNNs as a computational model [25]. The convolutional neural networks have been subject to other studies [72]. In a GCN model, the vertices are updated by aggregation of the information from the neighbouring vertices by considering a relational inductive bias. Here, the spacial-based GCN model is implemented that applies convolution spacial domain based on the initial graph structure, mathematically modelled in Ref. [25]. A nonlinear activation function *f* (such as a sigmoid or softmax) applies at each layer for each vertex *u* with parameters (**b**_l_, **Γ**_l_) (for *l* = 0, ⋯, *L*), where the inputs are the vertex embeddings and adjacency matrix. The activation function applies to every vertex embedding as [25]:

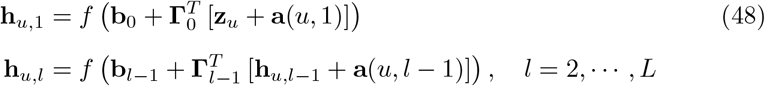

where **a**(⋯) is the aggregation function:

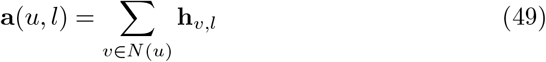

where *N*(*u*) is the list of the neighbouring vertices of *u*. The outputs are again predicted as follows:

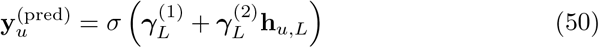

In our training, we considered a dataset of *T* graphs *G*_t_(*V, E*, **A, P**) and the measurements **Y**^(t)^ (*t* = 0, 1, ⋯,*T* − 1). The dataset is randomly split into training dataset (80 %) *T*_train_, validation dataset (10 %) *T*_valid_, and test dataset (10 %) *T*_test_. The test dataset is such that it does not share any information with training or validation datasets. In this way, the set of training parameters consists of {**b**_l_, **Γ**_l_} for *l* = 0, 1, ⋯, *L*. We used the so-called *Adam* algorithm and the MSE loss function as an optimiser. The mini-batches are chosen as a random subset of graph vertices at each training iteration. Other aggregation functions are also chosen for testing, such as the mean or maximum pooling aggregation functions:

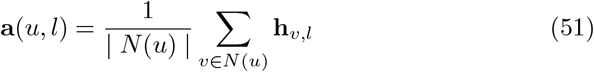

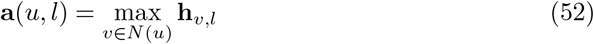

A jump knowledge connection algorithm [73] was used, according to which the model will additionally apply a final linear transformation to transform vertex embeddings to the expected output feature dimensionality. The long shortterm memory (lstm) algorithm [74] was used. Furthermore, a DeeperGCNLayer model [75] was implemented, including the pre-activation residual connection, the residual connection, the dense connection, and no connections.

#### 4.6.3. Test Case

The first test system is an intuitive system of five vertices, with a local transfer entropy given as:

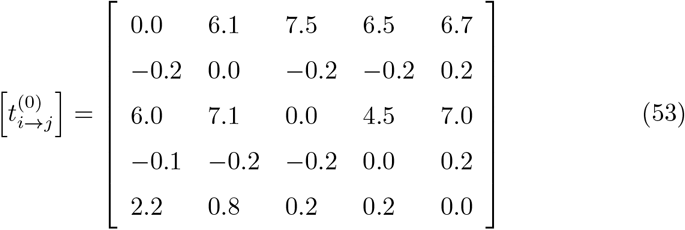

At every time iteration, a Gaussian-like noise 𝒩 (0, 1) (mean value zero and standard deviation one) is added to every element of the matrix as:

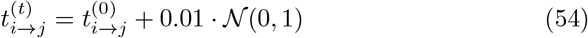

It is easy to identify which vertex 1 has the highest rank and which vertex 3 has the second highest. The length of the dataset is *T* = 2000 frames, and the lengths of the training, validation, and test datasets are 80 %, 10 %, and 10 %, respectively. The batch sizes are (100, 1, 1) and several epochs of 250 iterations on the training dataset. In this case, six layers with 32 channels, each layer is used as a setup model. The *adam* optimiser is chosen, the activation function for the hidden layers is the so-called *relu*, and the sigmoid function for the output layer. The aggregation function is the so-called *maximum*, and the jumping knowledge mode is the so-called *lstm*. The learning rate parameter is fixed at a value of 0.001.

Figure 15(a) shows the loss as a function of epochs for the training dataset, and Fig. 15(b) shows the performance metrics (such as mean absolute error (mae) and root mean square error (rmse)) for the training, validation, and test datasets. The 95 % confidence intervals for the means are also shown in each plot, calculated using the bootstrapping algorithm [76]. Our results indicate that only a few epochs (typically, 50 epochs) are needed to optimise the parameters of the machine learning model.

**Figure 15:**
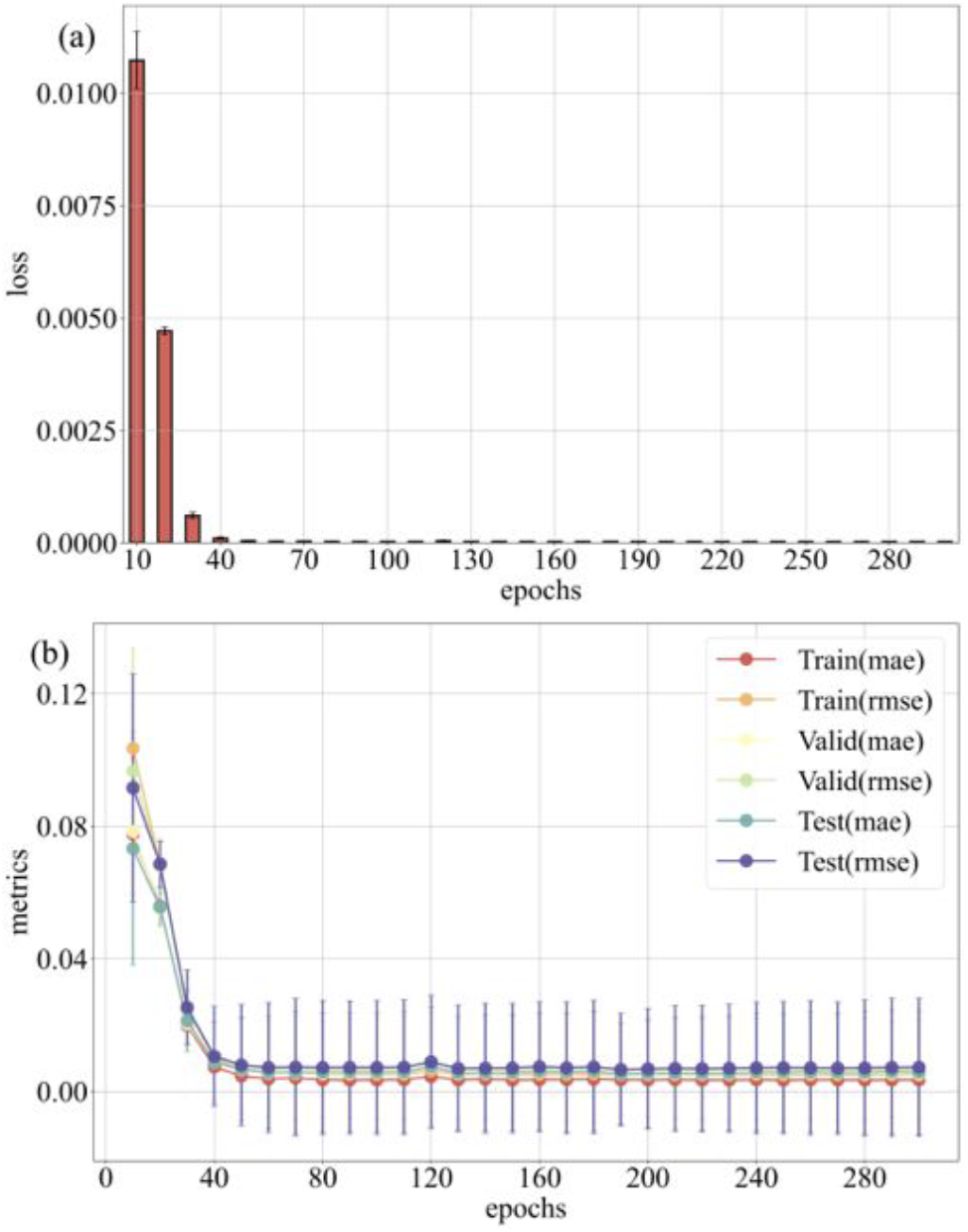
Test system: (a) Loss as a function of epochs for the training dataset; (b) Performance metrics (mean-absolute-error and root-mean-square-error) for the training, validation, and test datasets.

Furthermore, in Fig. 16(a)-(c), we present the predicted and the actual PageRank distributions of the training, validation, and test datasets. The results indicate excellent agreement between the predicted and true distributions for the three datasets.

**Figure 16:**
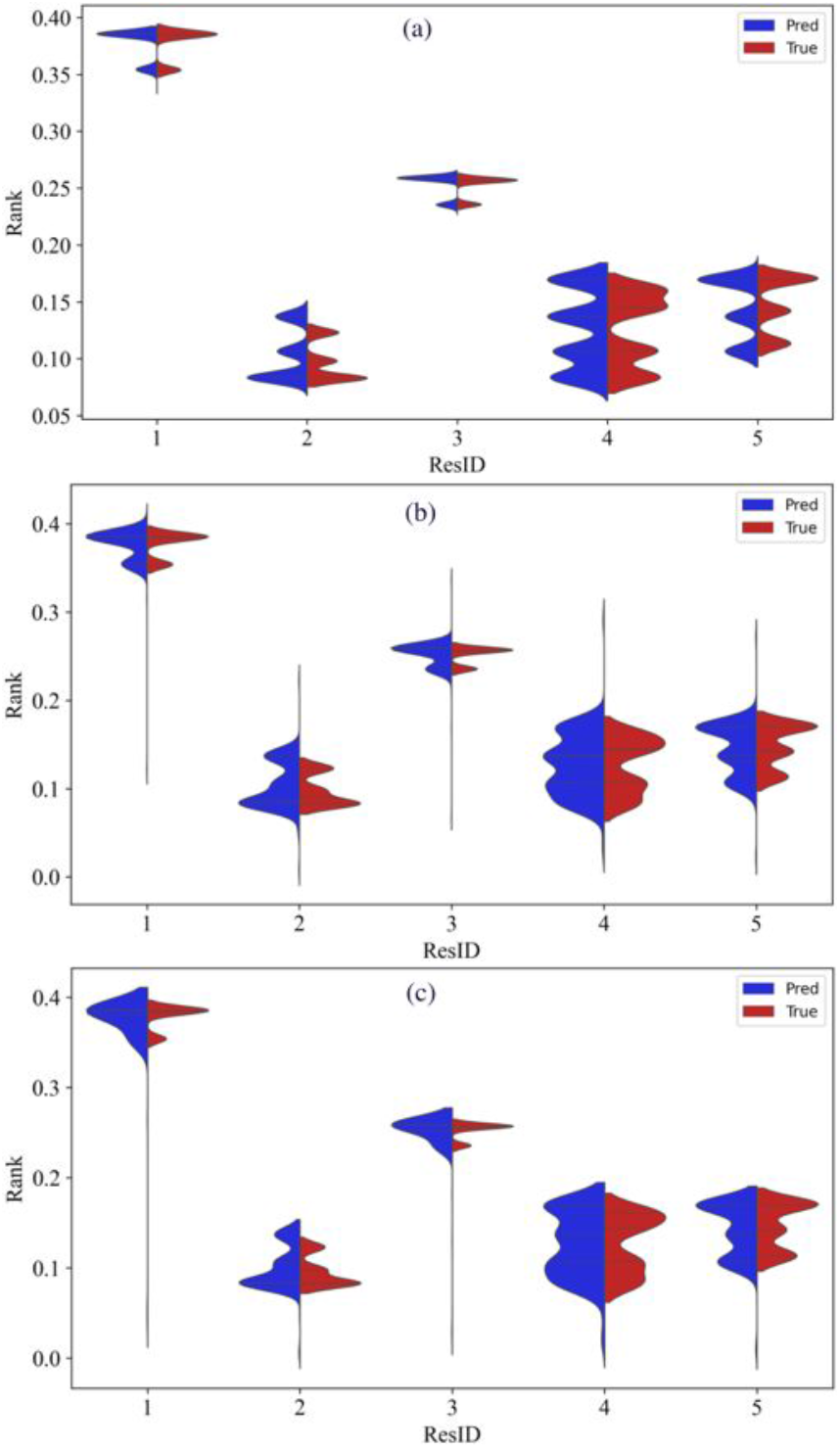
Test system: Predicted and the true (measured) PageRank distributions based on (a) the training dataset, (b) the validation dataset, and (c) the test dataset.

#### 4.6.4. BPTI[14-38]Abu Protein

Next, we considered the biological system, BPTI[14-38]Abu protein. In this study, the Tsallis local transfer entropy matrix 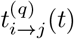 between the amino acids *I* and *j* is obtained from short MD simulations data (*t* = 0, 1, ⋯,*T* −1). We used the kernel density estimator approach with *q* = 0.85, as described in Supporting Information. The dataset size is *T* = 1000 frames, and the training, validation, and test dataset lengths are 80 %, 10 %, and 10 %, respectively. The batch sizes are (100, 1, 1) and several epochs of 1000. Here, the number of layers is ten, and the number of channels at each layer is 16. Again, the *adam* optimiser is used with a *relu* activation function for the hidden layers and the sigmoid function for the output layer. The aggregation function again was *maximum* and the *lstm* jumping knowledge mode. The learning rate parameter is 0.001.

Figures 17(a)-(b) present the loss as a function of epochs for the training dataset and the performance metrics for the training, validation, and test datasets. Again, the results show that in about 50-100 epochs the parameters of the computational model are optimised.

**Figure 17:**
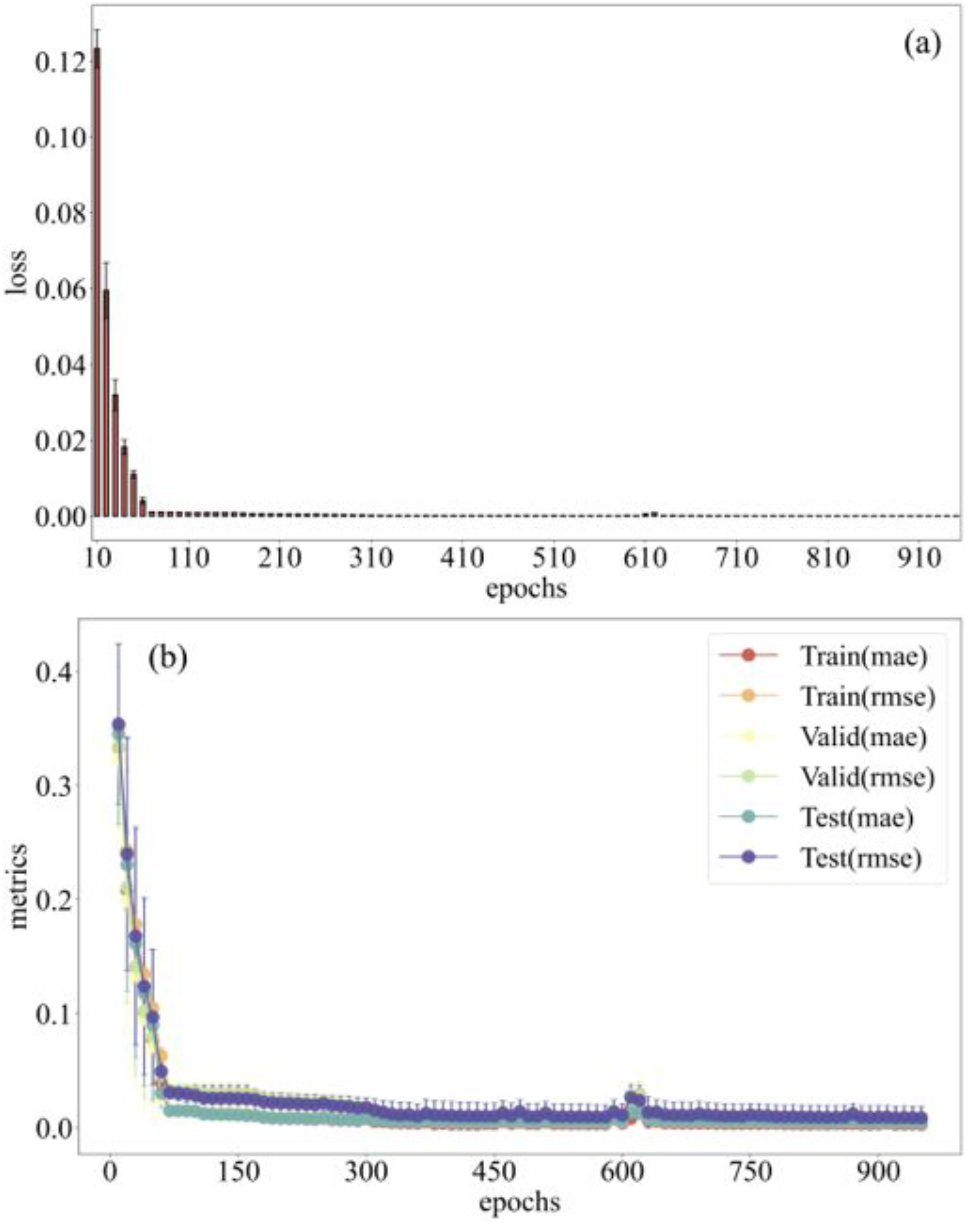
BPTI[14-38]Abu protein: (a) Loss as a function of epochs for the training dataset; (b) Performance metrics (mean-absolute-error and root-mean-square-error) for the training, validation, and test datasets.

In Fig. 18(a)-(c), we also present the predicted and the measured (true) PageRank values based on the training, validation, and test datasets, which again show good agreement between the predicted and true distributions in all datasets.

**Figure 18:**
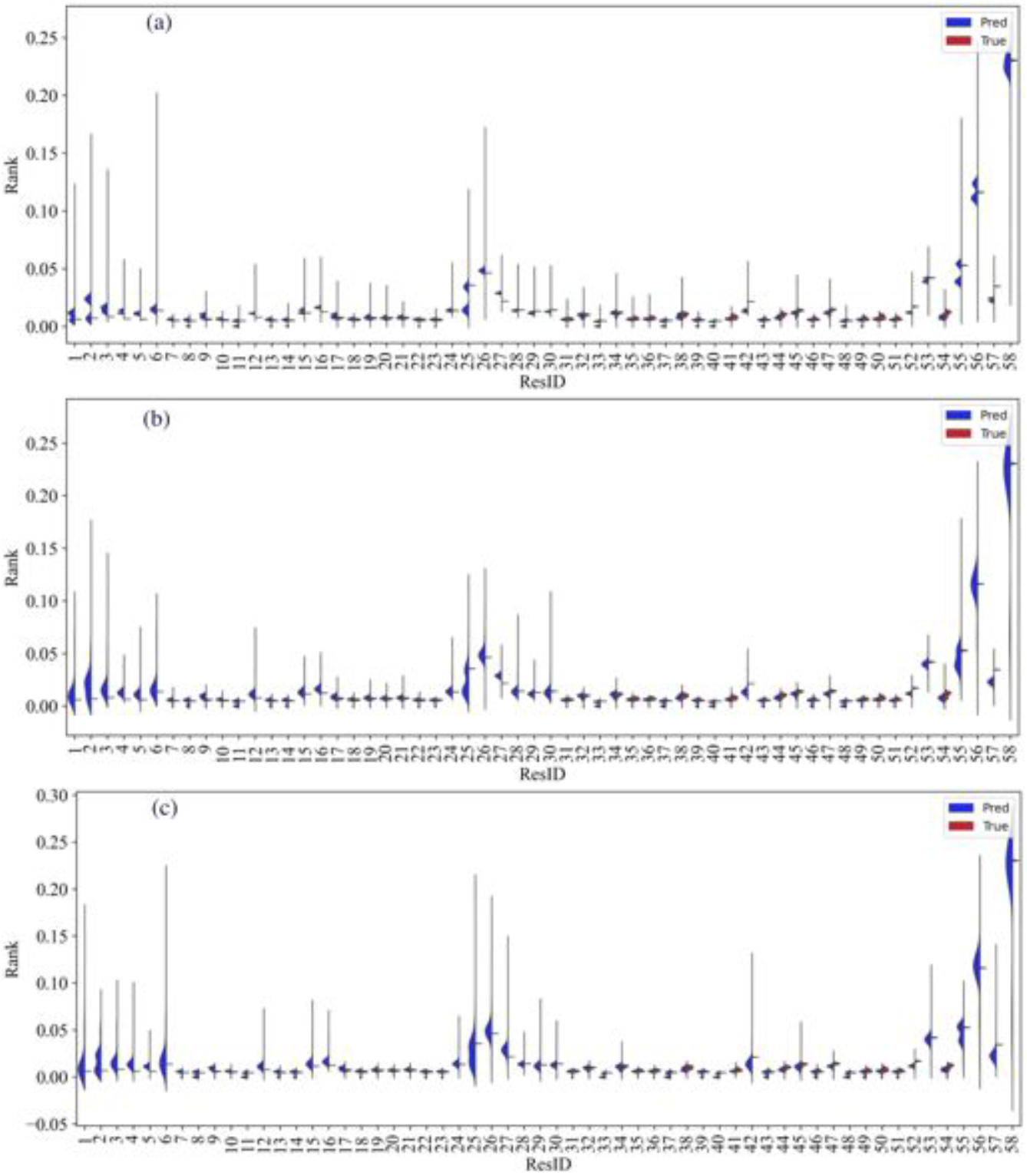
BPTI[14-38]Abu protein:: Predicted and the true (measured) PageRank distributions based on (a) the training dataset; (b) the validation dataset; and (c) the test dataset.

Our results indicate that the DGCNN model can predict the distributions of the PageRank values for all datasets, including the test dataset. Note that computations of the local transfer entropies can be costly; therefore, developing predictive machine learning models of the PageRank distributions relying on short length measurements can reduce significantly the computation demands.

## 5. Conclusions

This study discussed some alternative definitions for the information-theoretic measures of transfer entropy between time series using (*α, q*)-framework measures to validate statistical efficiency. The primary goal was to develop new measures of high-order correlations that preserved information-theoretic properties of information transfer due to local operations.

Besides, this study revealed the decoded information contained in an amino acid sequence establishing a 3D protein structure by comparing the physical-chemical properties with the ranked role in the protein interaction network topology using new graph-theoretic measures based on the constructed DiGraph models of (*α, q*) transfer entropy.

Moreover, in this study, the DGCNN model was used to classify the amino acid influence in a protein structure trained upon short equilibrium structure fluctuations at sub-nanosecond time scales, as obtained from MD simulations. For that, a heat flow kernel DiGraph embedding theoretical framework was developed. In this machine learning computational model, the amino acid information (represented by the vertices of a directed graph) is encoded by an Euclidean vector space embedded vector. It may be interesting to explore the mapping into other vector spaces using the differential geometry of a graph structure.

We examined the influence of disulphide bridges on the 3D structure of the BPTI wild- and mutated-type proteins.

## Acknowledgments

The author is grateful to the International Balkan University supporting the research activity.

## Data Availability

The supplementary materials used in this study are available within the article and its Supporting Information. The molecular dynamics simulations data used in this study are available from the corresponding author upon request.

## Footnotes

1 biophysics.cs.vt.edu

2 www.charmm.org

