## Supplemental Figure 1 to 10 for "Elucidating Partial Folding State of Bovine Pancreatic Trypsin Inhibitor by a Combined Study of Molecular Dynamics Simulations, Information Theory, Molecular Graph Theory, and Machine Learning"


Hiqmet Kamberaj*

*International Balkan University, Department of Computer Engineering, Makedonsko-Kosovska Brigada BB, Skopje, R of North Macedonia*


---


**Abstract**

Using a notably large amount of data in investigating physical and chemical phenomena demands new statistical and computational approaches; besides, the cross-validations require well-established theoretical frameworks. This study aims to validate the statistical efficiency of alternative definitions for the information-theoretic measures, such as transfer entropy, using the so-called $(\alpha, q)$-framework. The primary goal is to find measurements of high-order correlations that preserve information-theoretic properties of information transfer between the components of a dynamical system (such as a protein) due to local operations. Besides, this study aims to decode the information contained in the amino acid sequence establishing a three-dimensional protein structure by comparing the amino acids physical-chemical properties with their ranked role in the protein interaction network topology using new graph-theoretic measures based on the constructed digraph models of $(\alpha, q)$ information transfer within a heat flow kernel embedding framework. Moreover, this study aims to use the Deep Graph Convolution Neural Networks for classifying the role of each



*Corresponding author

amino acid in a protein trained upon short equilibrium structure fluctuations at sub-nanosecond time scales.

In particular, this study examines the influence of disulphide bridges on the three-dimensional structure of the Bovine Pancreatic Trypsin Inhibitor wild type and mutated analogue protein.
*Keywords:* $(\alpha, q)$ Transfer Entropy, Heat Kernel Molecular DiGraph Embedding, Heat and Information Flow, Deep Graph Convolution Neural Networks, Gibbs Free Energy

### 1. The ($\alpha$, $q$) Information Theoretic Measures

#### *1.1. Markovian Processes*

The Granger causality [1] concept is usually used to characterise the dependence of one variable $Y$ measured over time on another variable $X$ measured synchronously. This concept is initially being used to define the direction of interaction by estimating the contribution of $X$ in predicting $Y$. However, there exist many other variations of this concept, such as the linear approaches in the time and frequency domain.

The information theory measure of transfer entropy quantifies the statistical coherence between two processes that evolve in time. Schreiber [2] introduced the transfer entropy as the deviation from the independence of the state transition (from the previous state to the next state) of an information destination $X$ from the (previous) state of an information source $Y$.

We will use two vectors to characterise the dynamics of the two random Markovian processes. For the first process, the vector is $\mathbf{X}^{\mu_x} = \{\mathbf{x}_k^{\mu_x}\}_{k=k_{0x}}^{T-1}$ where

$$\mathbf{x}_k^{\mu_x} = \langle x_{k-(m_x-1)\tau_x}, x_{k-(m_x-1)\tau_x+\tau_x}, \cdots, x_{k-\tau_x}, x_k \rangle$$

Here, $k_{0x} = (m_x - 1)\tau_x$, and the dynamics of the second process is characterised by the vector $\mathbf{Y}^{\mu_y} = \{\mathbf{y}_k^{\mu_y}\}_{k=k_{0y}}^{T-1}$:

$$\mathbf{y}_k^{\mu_y} = \langle y_{k-(m_y-1)\tau_y}, y_{k-(m_y-1)\tau_y+\tau_y}, \cdots, y_{k-\tau_y}, y_k \rangle$$

where $k_{0y} = (m_y - 1)\tau_y$. In the following discussion, $\mathbf{x}_k^{\mu_x+1}$ represents the vector

$$\mathbf{x}_k^{\mu_x+1} = \langle x_{k-(m_x-1)\tau_x}, x_{k-(m_x-1)\tau_x+\tau_x}, \cdots, x_{k-\tau_x}, x_k, x_{k+\delta} \rangle$$

Similarly, $\mathbf{y}_k^{\mu_y+1}$ represents the vector

$$\mathbf{y}_k^{\mu_y+1} = \langle y_{k-(m_y-1)\tau_y}, y_{k-(m_y-1)\tau_y+\tau_y}, \cdots, y_{k-\tau_y}, y_k, y_{k+\delta} \rangle$$

Note that $m$ and $\tau$ are characteristics of each random process. The choice of $m$ and $\tau$ is crucial in order to reconstruct the dynamical structure of the random processes as discussed above. For clarity, we denote $\mu + 1 \equiv (m + 1, \tau)$ and $k_0 = \max\left((m_x - 1)\tau_x, (m_y - 1)\tau_y\right)$.

*1.2. Embedding Vector Analysis*

In this study, we employed an encoding approach to embed the vector states to a sequence of symbols. This encoding is an improved version of the previous studies [3, 4, 5, 6]. The idea is in finding an encoder or transformation $E$ such that

$$\langle i_1, i_2, \cdots, i_m \rangle = E\left(\langle x_1, x_2, \cdots, x_m \rangle\right) \tag{1}$$

where $\langle i_1, i_2, \cdots, i_m \rangle$ is the output symbols state vector and $\langle x_1, x_2, \cdots, x_m \rangle$ is the input state vector of the real numbers. Here, $E$ is the so-called *Encoder* or *Transformation* mapping. The improvement in comparison with other approaches [3, 4, 5, 6] consists in that here only one encoder, $E$, uniquely maps each state vector to a sequence of symbols. For that, we used the *class LabelEncoder* from the module *sklearn.preprocessing._label* of the Python programming language.

The pseudo-code is shown in the following:

```
from sklearn.preprocessing import LabelEncoder
import numpy as np
Symbols = ['a', 'b', 'c', 'd', 'e', 'f', 'g', 'h', 'i', 'j', 'k', 'l', 'm', 'n', 'o',
'p', 'q', 'r', 's', 't', 'u', 'v', 'w', 'x', 'y', 'z']
def embdSymbolEncoder(x, Nt=1000, M=3, Tau=1, delta=1, Nbins=100):
"""
Symbolic encoding function
"""
```

```
Ndata = Nt - (M-1)*Tau - delta
xmin = x.min()
xmax = x.max()
Nbins = int( Ndata**(1.0/M) ) + 1
if Nbins < 2: Nbins=2
if Nbins > 26: Nbins=26
S = np.linspace(xmin, xmax, Nbins)
le = LabelEncoder()
le.fit( S )
inds = np.digitize(x, S) - 1
Xs = list()
X1s = list()
X = np.zeros((M,), dtype=float)
X1 = np.zeros((M+1,), dtype=float)
for k in range((M-1)*Tau, Nt-delta):
m = np.arange(M)
X[:M] = S[inds[k - m*Tau]]
labels = le.transform(X)
Xs.append( [Symbols[label] for label in labels] )
X1[:M] = S[inds[k - m*Tau]]
X1[M:] = S[inds[k + delta]]
m = np.arange(M+1)
labels = le.transform(X1)
X1s.append( [Symbols[label] for label in labels] )
return Xs, X1s
```

#### *1.3. The α-Information Theoretic Measures*

The so-called Rényi entropy is defined as [7]:

$$H_\alpha(X) = \frac{1}{1-\alpha}\log\left(\sum_k \left(p(\mathbf{x}_k^{\mu_x})\right)^\alpha\right) \qquad (2)$$

which is non-additive from Penrose's viewpoint [8]. Here, $\log(\cdots)$ is the natural logarithm and thus the units are of *nats*. In Eq. 2, $\alpha$ is a real number parameter such that $\alpha \neq 1.0$. For $\alpha = 1.0$, the Shannon entropy is obtained [9]. In Fig. 1, a comparison of Shannon entropy ($\alpha = 1.0$) with Rényi entropies for different values of $\alpha$ is shown for an experiment of two possible outcomes: *success* (binary bit 1) with probability $p$ and *failure* (binary bit 0) with probability $1-p$. Here, $X$ follows the Bernoulli probability distribution with parameter $p$, $Ber(p)$; therefore,

$$H_\alpha(X) = \frac{1}{1-\alpha} \log\left(p^\alpha + (1-p)^\alpha\right) \tag{3}$$

$$H(X) = -\left[p \log p + (1-p)\log(1-p)\right] \tag{4}$$

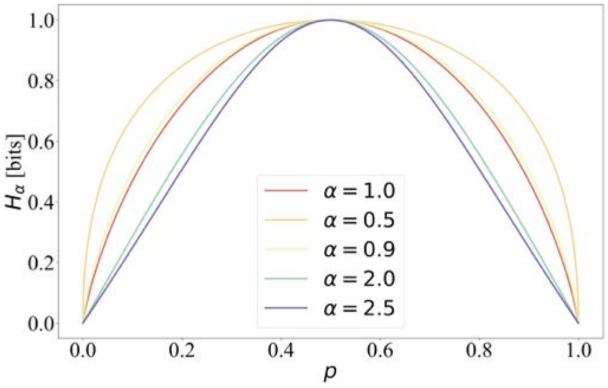


Figure 1: Comparison of Shannon ($\alpha = 1.0$) and Rényi entropies for different values of $\alpha$. The experiment has two possible outcomes following the Bernoulli probability distribution with parameter $p$, $Ber(p)$.

The so-called *Rényi mutual information* can be defined as [10]:

$$I_\alpha^{(1)}(X;Y) = H_\alpha(X) + H_\alpha(Y) - H_\alpha(X,Y) \tag{5}$$

Note that $I_\alpha^{(1)}(X;Y)$ can be negative, in particular for weakly coupled systems; therefore, $I_\alpha^{(1)}(X;Y)$ does not maximise the mutual information and the channel capacity, defined as [11]: $C = \max_{p(x)} I(X;Y)$ where the maximum is taken over all possible input distributions $p(x)$.

An alternative definition of the Rényi mutual information is given as [12] (and the references therein):

$$I_{\alpha}^{(2)}(X;Y) = H_{\alpha}(Y) - H_{\alpha}(Y|X) \tag{6}$$

which can be interpreted as an average information gain on the output $Y$ after an input symbol is received by the channel. In Eq. 6, $H_{\alpha}(Y|X)$ is the conditional Rényi entropy:

$$H_{\alpha}(Y|X) = \frac{\alpha}{1-\alpha} \log \sum_{k_x} p(\mathbf{x}_{k_x}^{\mu_x}) \left( \sum_{k_y} \left[ p(\mathbf{y}_{k_y}^{\mu_y} | \mathbf{x}_{k_x}^{\mu_x}) \right]^{\alpha} \right)^{\frac{1}{\alpha}} \tag{7}$$

Interestingly, $I_{\alpha}^{(2)}(X;Y)$ satisfies the properties of the information transfer [12] and $H_{\alpha}(Y|X) \leq H_{\alpha}(Y)$; thus, $I_{\alpha}^{(2)}(X;Y) \geq 0$. Furthermore, $I_{\alpha}^{(2)}(X;Y)$ maximises the mutual information and the channel capacity.

The joint Rényi entropy is defined as:

$$H_{\alpha}(X,Y) = \frac{1}{1-\alpha} \log \left( \sum_{k_x,k_y} \left( p(\mathbf{x}_{k_x}^{\mu_x}, \mathbf{y}_{k_y}^{\mu_y}) \right)^{\alpha} \right) \tag{8}$$

*1.4. The q-Framework Information Theoretic Measures*

The so-called Tsallis entropy is defined as [13]:

$$\begin{aligned} H_q(X) &= \frac{1}{1-q} \left( \sum_{k} (p(\mathbf{x}_k^{\mu_x}))^q - 1 \right) \\ &= \sum_{k} p(\mathbf{x}_k^{\mu_x}) \log_q \left( \frac{1}{p(\mathbf{x}_k^{\mu_x})} \right) \end{aligned} \tag{9}$$

where the function $\log_q(x)$ is given as [13, 14, 15]:

$$\log_q(x) = \frac{x^{1-q} - 1}{1-q} \tag{10}$$

and $q$ is a real number parameter, in general, $q \neq 1.0$. For $q = 1.0$, the Shannon entropy is obtained.

$H_q(X)$ is also non-additive, based on Penrose's definition [8] and furthermore, it belongs to the class of non-extensive statistical mechanics [15] (and the references therein). A comparison of Shannon entropy ($q = 1.0$) with Tsallis entropies for different values of $q$ is shown in Fig. 2. Again, for illustration, the experiment of two possible outcomes, *success* (binary bit 1) with probability $p$ and *failure* (binary bit 0) with probability $1 - p$, is chosen where $X$ follows the Bernoulli probability distribution with parameter $p$.

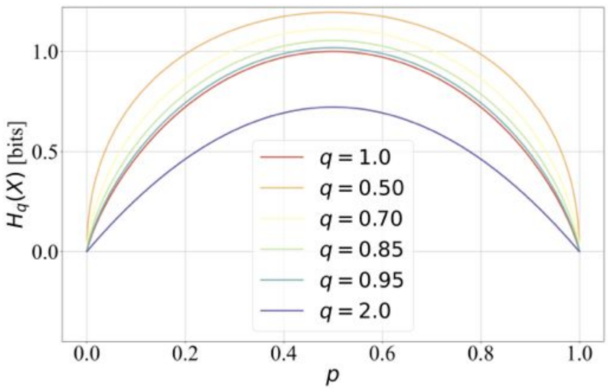


Figure 2: Comparison of Shannon ($q = 1.0$) and Tsallis entropies for different values of $q$. For illustration, the experiment of two possible outcomes is considered following the Bernoulli probability distribution with parameter $p$, $Ber(p)$.

The Tsallis statistics can be used to define measures of the information transfer [16], such as the so-called *Tsallis mutual information* [17]:

$$I_q^{(1)}(X;Y) = H_q(Y) - H_q(Y|X) = H_q(X) + H_q(Y) - H_q(X,Y) \tag{11}$$

$I_q^{(1)}(X;Y)$ can also be negative (such as for weakly coupled systems), and hence it does not maximise the mutual information and the channel capacity. In Eq. 11, $H_q(Y|X)$ is the conditional Tsallis entropy [17]:

$$\begin{aligned} H_q(Y|X) &= \frac{1}{1-q}\sum_{k_x} p(\mathbf{x}_{k_x}^{\mu_x})\left(\sum_{k_y}\left[p(\mathbf{y}_{k_y}^{\mu_y}|\mathbf{x}_{k_x}^{\mu_x})\right]^q - 1\right) \\ &= \sum_{k_x,k_y} p(\mathbf{x}_{k_x}^{\mu_x}, \mathbf{y}_{k_y}^{\mu_y}) \log_q\left(\frac{1}{p(\mathbf{y}_{k_y}^{\mu_y}|\mathbf{x}_{k}^{\mu_x})}\right) \end{aligned} \tag{12}$$

such that $H_q(X,Y) = H_q(X) + H_q(Y|X)$ [17].

Another definition of the Tsallis mutual information is given, taking into account the nonextensivity correction term:

$$I_q^{(2)}(X;Y) = H_q(X) \oplus_q H_q(Y) - H_q(X,Y) \tag{13}$$

where $\oplus_q$ function is given as [18]:

$$x \oplus_q y = x + y + (1-q)xy \tag{14}$$

Note that the nonextensive correction term, $(1-q)H_q(X)H_q(Y)$ (for $q < 1$), may reduce the negativity of $I_q^{(2)}(X;Y)$ compared with $I_q^{(1)}(X;Y)$.

Moreover, we can define the so-called *Tsallis divergence*, based on $q$-framework, as [17, 18]:

$$I_q^{(3)}(X;Y) = \sum_{k_x,k_y} p(\mathbf{x}_{k_x}^{\mu_x}, \mathbf{y}_{k_y}^{\mu_y}) \log_q \frac{p(\mathbf{x}_{k_x}^{\mu_x}, \mathbf{y}_{k_y}^{\mu_y})}{p(\mathbf{x}_{k_x}^{\mu_x})p(\mathbf{y}_{k_y}^{\mu_y})} \tag{15}$$

*1.5. $(\alpha, q)$-Framework Mutual Information*

The so-called *Sharma-Mittal entropy* is defined as [19, 12]:

$$H_{\alpha,q}(X) = \frac{1}{1-q}\left(\left(\sum_k \left(p(\mathbf{x}_k^{\mu_x})\right)^{\alpha}\right)^{\dfrac{1-q}{1-\alpha}} - 1\right) \tag{16}$$

which can be expresses as [12]

$$H_{\alpha,q}(X) = \eta_q(H_\alpha(X)) \tag{17}$$

where

$$\eta_q(x) = \frac{\exp\left((1-q)x\right) - 1}{1-q} \tag{18}$$

and $H_\alpha(X)$ is the Rényi entropy given by Eq. 2. Here, $(\alpha, q)$ are two real valued parameters (in general, $\alpha \neq 1$ and $q \neq 1$), as defined for the Rényi and Tsallis frameworks, respectively. Furthermore, the conditional Sharma-Mittal entropy relates to the conditional Rényi entropy $H_\alpha(Y|X)$ (see Eq. 7) as:

$$H_{\alpha,q}(Y|X) = \eta_q(H_\alpha(Y|X)) \tag{19}$$

Note that for $q = 1$ and $\alpha \neq 1$, the Sharma-Mittal equals the Rényi entropy and, for $\alpha = q \neq 1$, it equals the Tsallis entropy. When $\alpha = q = 1.0$, the Shannon entropy is obtained.

A comparison of Shannon entropy ($\alpha = 1.0$, $q = 1.0$) with Sharma-Mittal entropies [19] for different values of ($\alpha$, $q$) is shown in Fig. 3. Again, the experiment of two possible outcomes, *success* (binary bit 1) with probability $p$ and *failure* (binary bit 0) with probability $1 - p$, is chosen where $X$ follows $Ber(p)$.

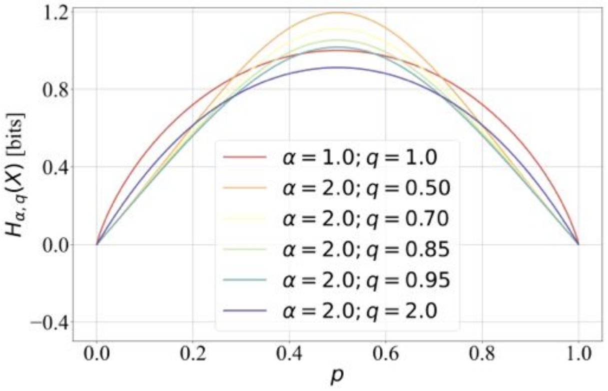


Figure 3: Comparison of Shannon ($\alpha = 1.0$, $q = 1.0$) and Sharma-Mittal entropies for different values of ($\alpha$, $q$). As an example, the experiment of two possible outputs is considered following the Bernoulli probability distribution with parameter $p$, $Ber(p)$.

The so-called $(\alpha, q)$ information theoretic measures have already been formulated by Ilić and Djordjević [12]. In particular, the $(\alpha, q)$ mutual information and $(\alpha, q)$ capacities have been introduced, which are briefly summarised in the following, as in Ref. [12].

The first definition of $(\alpha, q)$ mutual information can be as

$$I^{(1)}_{\alpha,q}(X;Y) = H_{\alpha,q}(X) + H_{\alpha,q}(Y) - H_{\alpha,q}(X,Y) \tag{20}$$

Note that $I^{(1)}_{\alpha,q}(X;Y)$ can also be negative, in particular for weakly coupled systems [10].

A second definition [12] that satisfies the properties of the information transfer is given:

$$I^{(2)}_{\alpha,q}(X;Y) = H_{\alpha,q}(Y) \ominus_q H_{\alpha,q}(Y|X) = \eta_q(I^{(2)}_{\alpha}(X;Y)) \tag{21}$$

where

$$x \ominus_q y = \frac{x - y}{1 + (1-q)y} \tag{22}$$

Using Eq. 21, an alternative definition of Tsallis mutual information can also be introduced for $\alpha = q \neq 1$:

$$I^{(4)}_{q}(X;Y) = H_q(Y) \ominus_q H_q(Y|X) = \eta_q(I^{(2)}_{\alpha=q}(X;Y)) \tag{23}$$

where

$$\begin{aligned} H_q(Y) &= H_{\alpha=q,q}(Y) \\ H_q(Y|X) &= H_{\alpha=q,q}(Y|X) \end{aligned} \tag{24}$$

and $I^{(2)}_{\alpha=q}(X;Y)$ is the Rényi mutual information $I^{(2)}_{\alpha}(X;Y)$, given by Eq. 6, for $\alpha = q$.

#### *1.6. Kernel Density Estimators*

In this study, we also employed kernel density estimators (KDEs) to approximate probability density functions. Shannon differential entropy $h(X)$ of a continuous random variable $X$ with density $f(x)$ is defined as

$$h(X) = -\int_{\Omega} f(x)\log f(x)\,dx = E[-\log f(X)] \tag{25}$$

where $\Omega$ is the support set of the random variable $X$ and $E[\cdots]$ denotes the expected value. Rényi differential entropy can be defined as:

$$h_\alpha(X) = \frac{1}{1-\alpha} \log \left( \int_\Omega (f(x))^\alpha \, dx \right) \tag{26}$$

where $f(x)$ is the probability density function of the continuous random variable $X$ and the integration is over the sample space $\Omega$ of $X$.

Tsallis differential entropy can also be defined as:

$$\begin{aligned} h_q(X) &= \frac{1}{1-q} \left( \int_\Omega (f(x))^q \, dx - 1 \right) \\ &= \int_\Omega f(x) \log_q \left( \frac{1}{f(x)} \right) dx = E \left[ \log_q \left( \frac{1}{f(X)} \right) \right] \end{aligned} \tag{27}$$

where $f(x) \neq 0$ for every $x \in \Omega$. Similarly, Sharma-Mittal differential entropy is calculated as

$$h_{\alpha,q}(X) = \eta_q(h_\alpha(X)) \tag{28}$$

Suppose we divide $X$ into histogram bins of length $\Delta$. The position of the centre of the histogram bin is

$$i\Delta \leq x_i < (i+1)\Delta, \quad i = -\infty, \cdots, -2, -1, 0, 1, 2, \cdots, +\infty$$

which is such that probability mass function of $x_i$ is

$$p(x_i) = f(x_i)\Delta = \int_{i\Delta}^{(i+1)\Delta} f(x) \, dx$$

Let $X^\Delta$ be the quantised random variable of $X$: $X^\Delta = x_i$. Then, it can be shown [11], if the density $f(x)$ of a continuous random variable $X$ is Riemann integrable:

$$\lim_{\Delta \to 0} \left( H(X^\Delta) + \log \Delta \right) = h(X) \tag{29}$$

If we express in units of bits, then the Shannon entropy $H(X^\Delta)$ of $n$-bit quantisation (that is, $\Delta = \frac{1}{2^n}$) of a continuous random variable $X$ is approximately:

$$H(X^\Delta) \approx h(X) + n \tag{30}$$

which indicates the number of bits required on the average to describe $X$ to $n$-bits accuracy is $n$.

For the Rényi entropy:

$$\begin{aligned} H_\alpha(X^\Delta) &= \frac{1}{1-\alpha}\log\left(\sum_i (p(x_i))^\alpha\right) \qquad (31) \\ &= \frac{1}{1-\alpha}\left[\log\left(\sum_i (f(x_i))^\alpha \Delta\right) + (\alpha-1)\log\Delta\right] \end{aligned}$$

which in the limit of infinitesimally small $\Delta$ gives:

$$\lim_{\Delta\to 0}\left(H(X^\Delta) + \log\Delta\right) = h_\alpha(X) \qquad (32)$$

Similarly, in bits ($\log \equiv \log_2$), the Rényi entropy $H_\alpha(X^\Delta)$ of $n$-bit quantisation, $\Delta = \frac{1}{2^n}$, of a continuous random variable $X$ is approximately:

$$H_\alpha(X^\Delta) \approx h_\alpha(X) + n \qquad (33)$$

which indicates the number of bits required on the average to describe $X$ to $n$-bits accuracy is $n$.

In the case of Tsallis entropy:

$$\begin{aligned} H_q(X^\Delta) &= \frac{1}{1-q}\left[\sum_i (p(x_i))^q - 1\right] \qquad (34) \\ &= \frac{1}{1-q}\left[\sum_i (f(x_i))^q \Delta^q - 1\right] \\ &= \frac{1}{1-q}\left[\sum_i (f(x_i))^q \Delta - 1\right]\Delta^{q-1} + \frac{\Delta^{q-1}-1}{1-q} \\ &= \frac{1}{1-q}\left[\sum_i (f(x_i))^q \Delta - 1\right]\Delta^{q-1} + \frac{\left(\frac{1}{\Delta}\right)^{1-q} - 1}{1-q} \\ &= \frac{1}{1-q}\left[\sum_i (f(x_i))^q \Delta - 1\right]\Delta^{q-1} + \log_q\left(\frac{1}{\Delta}\right) \end{aligned}$$

In the limit of infinitesimally small $\Delta$:

$$\lim_{\Delta\to 0}\left(H_q(X^\Delta) + \Delta^{q-1}\log_q\Delta\right) = h_q(X)\Delta^{q-1} \qquad (35)$$

When taking the limit $\Delta \to 0$, one has to distinguish two different sets of the values of $q$, $q < 1$ and $q > 1$. Figures 4(a)-(b) show the functions $\Delta^{q-1} \log_q \Delta$ and $\Delta^{q-1}$ plots for different values of $\Delta \to 0$: Fig. 4(a) for $q = 0.8$ and Fig. 4(b) $q = 2.5$. It can be seen that both these functions diverge to opposite signs as $\Delta \to 0$ for $q < 1$. On the other hand, for $q > 1$, $\Delta^{q-1}$ converges to zero and $\Delta^{q-1} \log_q \Delta$ and $\Delta^{q-1}$ to a constant value of $1/(1-q)$. Therefore, from Eq. 35, one can get:

$$H_q(X^\Delta) \approx \begin{cases} \Delta^{q-1}\left[h_q(X) - \log_q \Delta\right], & q < 1 \\ \dfrac{1}{q-1}, & q > 1 \\ h(X) - \log \Delta, & q = 1 \end{cases} \tag{36}$$

The values of $q < 1$ are practical to use because $h_q(X)$ can have positive and negative values, and thus $\Delta^{q-1}\left[h_q(X) - \log_q \Delta\right]$ can fluctuate around a finite value; in contrary, for $q > 1$, $H_q(X^\Delta)$ converges to a finite constant values for any random process $X$, and so it is not of practical interest because it does not maximise the mutual information and channel capacity.

For a discrete random variable $X$, the Sharma-Mittal entropy can be calculated as [19, 12]:

$$\begin{aligned} H_{\alpha,q}(X) &= \eta_q(H_\alpha(X)) \\ &= \frac{1}{1-q}\left[\left(\sum_i (p(x_i))^\alpha\right)^{\frac{1-q}{1-\alpha}} - 1\right] \end{aligned} \tag{37}$$

where $p(x_i)$ is the probability mass function at $x_i$. For a continuous random variable $X$, the Sharma-Mittal differential entropy can be calculated as

$$\begin{aligned} h_{\alpha,q}(X) &= \eta_q(h_\alpha(X)) \\ &= \frac{1}{1-q}\left[\left(\int_\Omega (f(x))^\alpha \, dx\right)^{\frac{1-q}{1-\alpha}} - 1\right] \end{aligned} \tag{38}$$

For a quantisation $X^\Delta$ of the continuous random variable of $X$: $X^\Delta = x_i$,

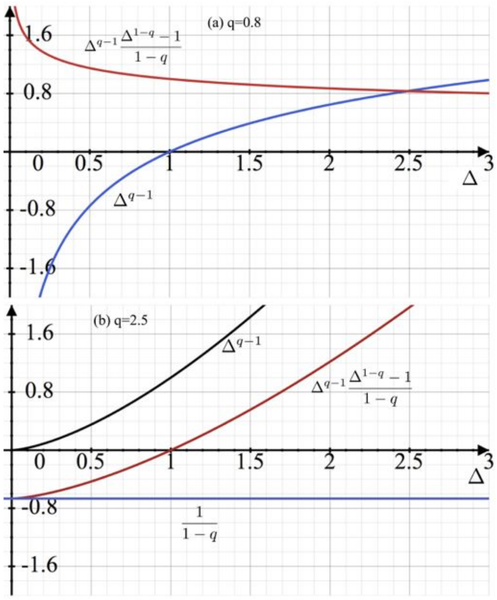


Figure 4: The functions $\Delta^{q-1}\log_q \Delta$ and $\Delta^{q-1}$ plots for different values of $\Delta \to 0$: (a) for $q = 0.8$ and (b) $q = 2.5$.

we have

$$
\begin{aligned}
H_{\alpha,q}(X^{\Delta}) &= \frac{1}{1-q}\left[\left(\sum_i (p(x_i))^{\alpha}\right)^{\dfrac{1-q}{1-\alpha}} - 1\right] \\
&= \frac{1}{1-q}\left[\left(\sum_i (f(x_i)\Delta)^{\alpha}\right)^{\dfrac{1-q}{1-\alpha}} - 1\right] \\
&= \frac{1}{1-q}\left[\left(\sum_i (f(x_i))^{\alpha}\Delta\right)^{\dfrac{1-q}{1-\alpha}} \Delta^{q-1} - 1\right] \\
&= \frac{1}{1-q}\left[\left(\sum_i (f(x_i))^{\alpha}\Delta\right)^{\dfrac{1-q}{1-\alpha}} - 1\right]\Delta^{q-1} + \frac{\left(\dfrac{1}{\Delta}\right)^{1-q} - 1}{1-q}
\end{aligned}
\tag{39}
$$

$$= \frac{1}{1-q}\left[\left(\sum_i (f(x_i))^\alpha \Delta\right)^{\dfrac{1-q}{1-\alpha}} - 1\right]\Delta^{q-1} + \log_q\left(\frac{1}{\Delta}\right)$$

In the limit $\Delta \to 0$:

$$\lim_{\Delta\to 0}\left(H_{\alpha,q}(X^\Delta) + \Delta^{q-1}\log_q \Delta\right) = h_{\alpha,q}(X)\Delta^{q-1} \tag{40}$$

Using the same argument as in the case of the Tsallis framework, we can write:

$$H_{\alpha,q}(X^\Delta) \approx \begin{cases} \Delta^{q-1}\left[h_{\alpha,q}(X) - \log_q \Delta\right], & q < 1 \\ \dfrac{1}{q-1}, & q > 1 \\ h_\alpha(X) - \log \Delta, & q = 1 \end{cases} \tag{41}$$

Consider two continuous random variables $X$ and $Y$ and a quantisation $(\Delta_x, \Delta_y)$ such that $X^{\Delta_x} = x_i$ and $Y^{\Delta_y} = y_i$. For the Sharma-Mittal framework of entropy, one can write:

$$\begin{aligned} H_{\alpha,q}(X^{\Delta_x}) &\approx \Delta_x^{q-1}\left[h_{\alpha,q}(X) - \log_q \Delta_x\right] \\ H_{\alpha,q}(Y^{\Delta_y}) &\approx \Delta_y^{q-1}\left[h_{\alpha,q}(Y) - \log_q \Delta_y\right] \\ H_{\alpha,q}(X^{\Delta_x}, Y^{\Delta_y}) &\approx (\Delta_x\Delta_y)^{q-1}\left[h_{\alpha,q}(X,Y) - \log_q(\Delta_x\Delta_y)\right] \end{aligned} \tag{42}$$

For $q > 1$, each of these terms converges to $1/(q-1)$, and thus

$$I_{\alpha,q}(X^{\Delta_x}; Y^{\Delta_y}) = H_{\alpha,q}(X^{\Delta_x}) + H_{\alpha,q}(Y^{\Delta_y}) - H_{\alpha,q}(X^{\Delta_x}, Y^{\Delta_y}) \to \frac{1}{q-1}$$

which does not maximise the mutual information and channel capacity; furthermore, the computed mutual information using finite resolution of the continuous random variables is not equal to that of continuous random variables. On the other hand, for $q = 1$ and $\alpha \neq 1$, we have

$$\begin{aligned} H_\alpha(X^{\Delta_x}) &\approx h_\alpha(X) - \log \Delta_x \\ H_\alpha(Y^{\Delta_y}) &\approx h_\alpha(Y) - \log \Delta_y \\ H_\alpha(X^{\Delta_x}, Y^{\Delta_y}) &\approx h_\alpha(X,Y) - \log(\Delta_x\Delta_y) \end{aligned} \tag{43}$$

Therefore,

$$I_\alpha(X^{\Delta_x}; Y^{\Delta_y}) = H_\alpha(X^{\Delta_x}) + H_\alpha(Y^{\Delta_y}) - H_\alpha(X^{\Delta_x}, Y^{\Delta_y})$$

$$\approx I_\alpha(X;Y)$$

which is approximately equal to $I_\alpha(X;Y)$ estimate of continuous random variables. Therefore, it can be suggested that for approximating the mutual information of the continuous random variables using KDE, the expression given by Eq. 21 is a better choice for every $\alpha$ and $q$:

$$\begin{aligned} I_{\alpha,q}(X^{\Delta_x};Y^{\Delta_y}) &= \eta_q\left(I_\alpha(X^{\Delta_x};Y^{\Delta_y})\right) \\ &\approx \eta_q\left(I_\alpha(X;Y)\right) \\ &= I_{\alpha,q}(X;Y) \end{aligned} \tag{44}$$

The expression given by Eq. 20 works better for every $\alpha$ and $q < 1$; however, in general, $I_{\alpha,q}(X^{\Delta_x};Y^{\Delta_y}) \neq I_{\alpha,q}(X;Y)$.

In this study, we used the following Python packages:
*from scipy import stats* from statistics,
*from sklearn.neighbors import KernelDensity* and *from sklearn.model_selection import GridSearchCV* from sklearn.
In statistics package, representation of a kernel-density estimate using Gaussian kernels was used. The class *gaussian_kde* works for both univariate and multivariate data. In particular, it includes automatic bandwidth determination. The estimation works best for a unimodal distribution; bimodal or multimodal distributions tend to be oversmoothed.

### 2. Linear and Non-linear Noisy Channels

For illustration, we considered two noisy channels (see also Fig. 5). The first channel is a XOR gate characterised by random Bernoulli distribution noise:

$$\begin{aligned} &X(t) = Ber(p), \quad \forall t \geq 0 \\ &Y(t) = X(t-1)\,\mathrm{XOR}\,Ber(p), \quad \forall t > 0 \end{aligned} \tag{45}$$

$$Y(0) = Ber(p)$$

where $Ber(p)$ is a random number generated at each iteration $t$ according to Bernoulli distribution with parameter $p$ and XOR function is such that:

$$a\,\mathrm{XOR}\,b = \begin{cases} 0, & \text{if } a = b \\ 1, & \text{otherwise} \end{cases} \tag{46}$$

Note that this is a non-linear binary channel; that is, $X$ and $Y$ dependence is highly non-linear. The second noisy channel is the following non binary valued linear channel:

$$\begin{aligned} X(t) &= A_x X(t-1) + \sigma_x \mathcal{N}(0,1) \\ Y(t) &= C_{xy} X(t-1) + \sigma_y \mathcal{N}(0,1), \quad \forall t > 0 \end{aligned} \tag{47}$$

where $A_x$, $C_{xy}$, $\sigma_x$, and $\sigma_y$ are parameters. $C_{xy}$ characterises the coupling strength between $X$ and $Y$ and $\mathcal{N}(0,1)$ is a random number following the standard normal distribution. Initially, $X(0) = \sigma_x \mathcal{N}(0,1)$ and $Y(0) = \sigma_y \mathcal{N}(0,1)$. Here, $\sigma_x$ and $\sigma_y$ characterise the strength of the external noise in the channel.

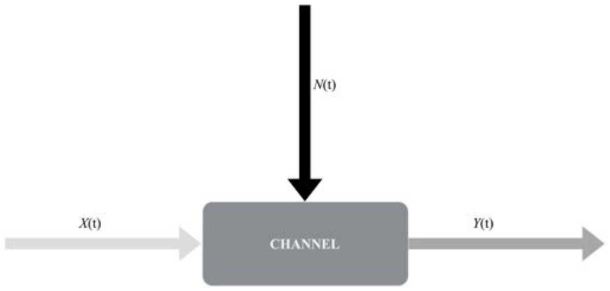


Figure 5: Graphical diagram of a noisy channel: $X(t)$ is the input signal at time $t$, $N(t)$ is the noise signal at $t$, and $Y(t)$ is the output signal at time $t$.

In Fig. 6, we show the XOR channel with different Bernoulli noise level by varying the parameter $p$. Fig. 6(a) presents the transfer entropy $T^{(\alpha,q)}_{X\to Y}$ in bits as a function of $p$ and Fig 6(b) the mutual information $I_{\alpha,q}(X;Y)$ in bits. We fixed $\alpha = 2.0$ and $q = 0.95$. Only the part for $p \in (0, 0.5]$ is shown because of the symmetry in $p$. Our results support the previous findings [12] that the Sharma-Mittal formulation of the entropy maximises the channel capacity ($C = \max_{p(X)} I(X;Y)$ [11]), in particular, for noisy and non-linearly coupled input-output signals.

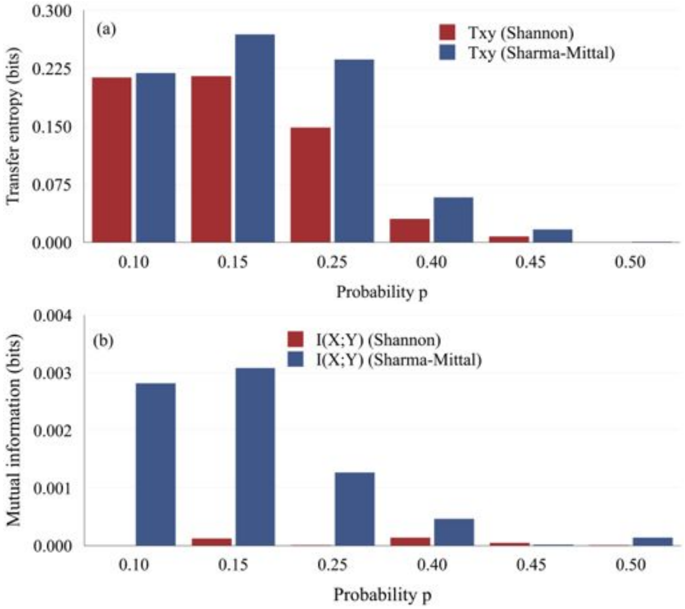


Figure 6: XOR channel with different Bernoulli noise level: (a) Transfer entropy $T^{(\alpha,q)}_{X\to Y}$ in bits for different Bernoulli distribution parameter $p$; (b) Mutual information $I_{\alpha,q}(X;Y)$ in bits. Here, $\alpha = 2.0$ and $q = 0.95$. Only the part for $p \in (0, 0.5]$ is shown because of the symmetry in $p$.

In Fig. 7, the linearly coupled Gaussian signals channel (as in Eq. 47) is presented with $A_x = 0.5$, $\sigma_x = \sigma_y = 0.1$. Besides, $\alpha$ and $q$ are shown in each plot of the computed transfer entropies and mutual informations versus coupling strengths $C_{xy}$ using Sharma-Mittal and Shannon entropies. Our results indicate that the Sharma-Mittal formulation maximises the transfer entropy while the mutual information is not significantly affected. Interestingly, these results prove the linearity dependence on input-output. (One can also look at the Refs. [20, 21] for some other related works.)

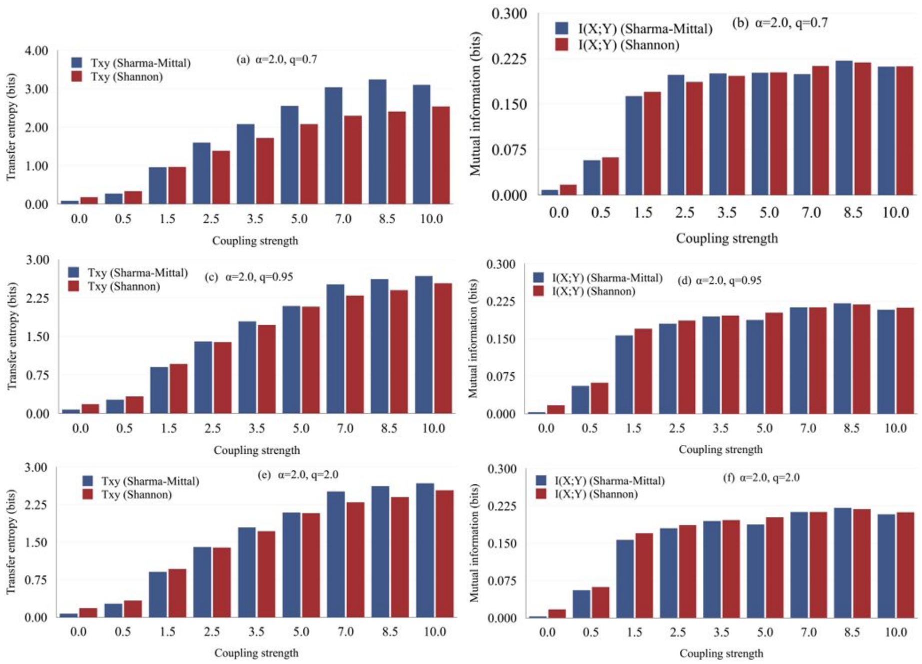


Figure 7: Linear Gaussian channel with $A_x = 0.5$, $\sigma_x = \sigma_y = 0.1$: (a), (c), and (e) Transfer entropy $T^{(\alpha,q)}_{X\to Y}$ in bits for different coupling strengths $C_{xy}$; (b), (d), and (f) Mutual information $I_{\alpha,q}(X;Y)$ in bits. $\alpha$ and $q$ are shown in each plot.

We also tested for higher strength of the Gaussian noise: $\sigma_x = \sigma_y = 1.0$, as shown in Fig. 8, for $A_x = 0.5$. Besides, $\alpha$ and $q$ are shown in each plot. The transfer entropies and mutual informations in bits versus coupling strengths $C_{xy}$ using Sharma-Mittal and Shannon entropies are presented. Again, our results indicate that for the linearly coupled signals the Sharma-Mittal formulation maximises the transfer entropy while the mutual information is not significantly affected for a broad range of $\alpha$ and $q$. Furthermore, in all our results the increasing trend of the transfer entropy and mutual information as a function of coupling strength is observed (see also Fig. 7 and Fig. 8).

For linear Gaussian channel with $A_x = 0.5$, $\sigma_x = \sigma_y = 0.1$, we compared the encoding and KDE (statistical package) transfer entropy computations as

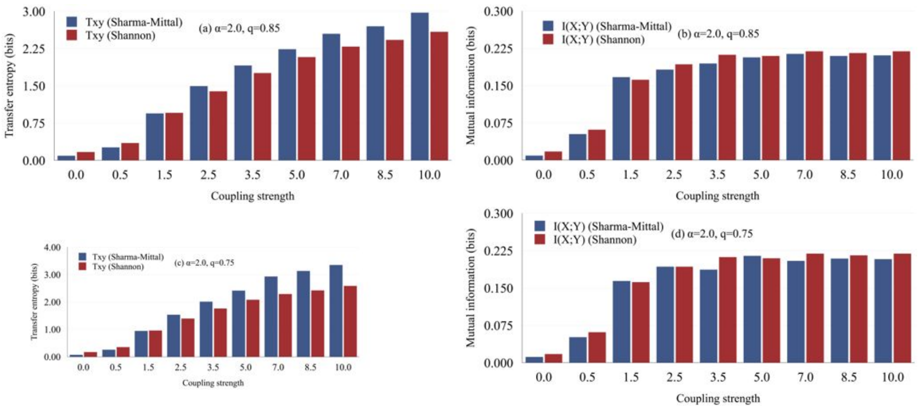


Figure 8: Linear Gaussian channel with $A_x = 0.5$, $\sigma_x = \sigma_y = 1.0$: (a) and (c) Transfer entropy $T^{(\alpha,q)}_{X\to Y}$ in bits for different coupling strengths $C_{xy}$; (b) and (d) Mutual information $I_{\alpha,q}(X;Y)$ in bits. $\alpha$ and $q$ are shown in each plot.

a function of coupling strength $C_{xy}$, as shown in Fig. 9(a) for Shannon entropy formulation and Fig. 9(b) for Sharma-Mittal entropy formulation with $\alpha = 2.0$ and $q = 0.85$.

In Fig. 10, we compare local transfer entropies calculated using the Shannon framework ($q = 1$) and Tsallis ($q = 0.85$) with symbolic encoding and KDE algorithms. The linear Gaussian channel with $A_x = 0.5$, $\sigma_x = 0.2$, $\sigma_y = 0.02$, and $C_{xy} = 12.5$ was considered. Note that there will be a difference (of about $\log\Delta$) between local transfer entropies calculated using symbolic encoding and KDE approach; however, in all calculations, one can see that the expected values are such that $t_{X\to Y} > 0$ and $t_{Y\to X}$ fluctuates around zero value.

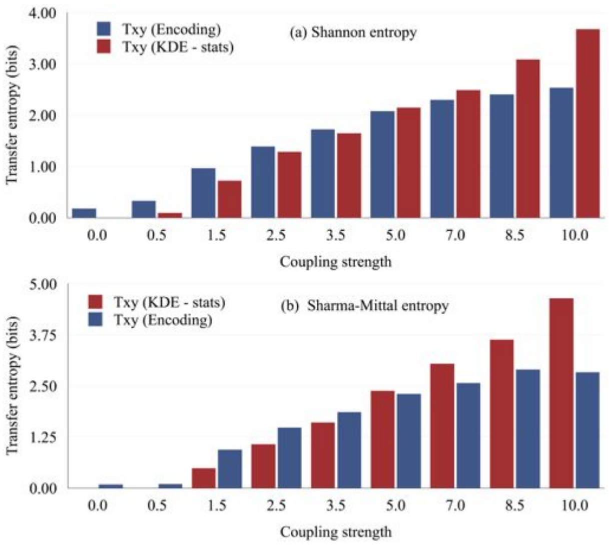


Figure 9: Linear Gaussian channel with $A_x = 0.5$, $\sigma_x = \sigma_y = 0.1$. Comparison of the encoding and KDE (statistical package) transfer entropy computations as a function of coupling strength $C_{xy}$: (a) Shannon entropy formulation; (b) Sharma-Mittal entropy formulation with $\alpha = 2.0$ and $q = 0.85$.

461–464.

[3] M. Staniek, K. Lehnertz, Symbolic transfer entropy, Phys. Rev. Lett. 100 (2008) 158101.

[4] H. Kamberaj, A. der Vaart, Extracting the causality of correlated motions from molecular dynamic simulations, Biophys. J. 97 (2009) 1747–1755.

[5] Dh. Nebiu, H. Kamberaj, Symbolic Information Flow Measurement (SIFM): A software for measurement of information flow using symbolic analysis, SoftwareX 11 (2020) 100470.

[6] H. Kamberaj, Heat flow random walks in biomolecular systems using symbolic transfer entropy and graph theory, J. Mol. Graph. Model. 104 (2021) 107838.

[7] A. Rényi, Contributions to the Theory of Statistics, in: Proceedings of the

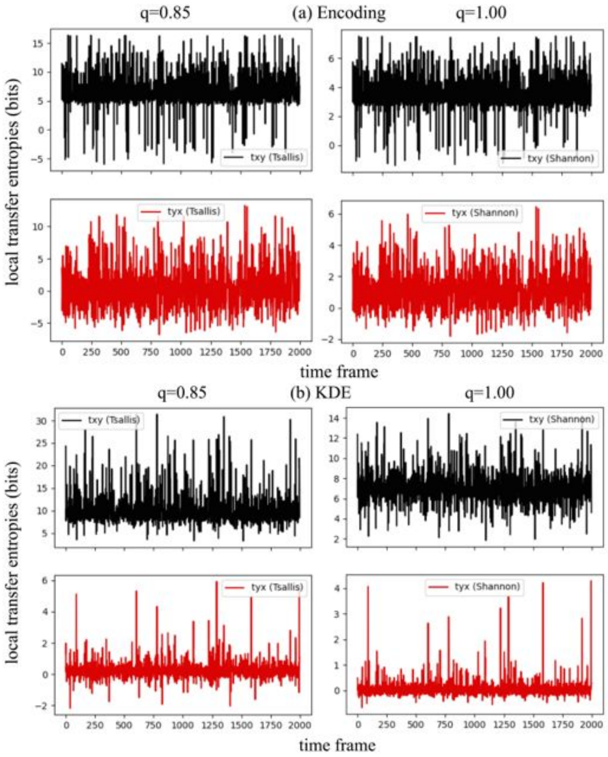


Figure 10: Linear Gaussian channel with $A_x = 0.5$, $\sigma_x = 0.2$, $\sigma_y = 0.02$, and $C_{xy} = 12.5$. Comparison of the encoding and KDE (statistical package) local transfer entropy computations as a function of time frame: (a) symbolic encoding algorithm; (b) KDE algorithm. Here, $q = 0.85$.

Fourth Berkeley Symposium on Mathematical Statistics and Probability, Vol. 1, University of California Press, 1961, pp. 547–561.

[8] O. Penrose, Foundations of Statistical Mechanics: A Deductive Treatment, Dover Publications, Mineola, NY, USA, 1970.

[9] S. Behrendt, T. Dimpfl, F. J. Peter, D. J. Zimmermann, RTransferEntropy − Quantifying information flow between different time series using effective transfer entropy, SoftwareX 10 (2019) 100265.

[10] S. O. Scalet, A. M. Alhambra, G. Styliaris, J. I. Cirac, Computable Rényi

mutual information: Area laws and correlations, Quantum 5 (2021) 541.

[11] M. C. Thomas, A. T. Joy, Elements of information theory, John Wiley & Sons, Inc., Hoboken, 2006.
